# Transcriptional pathways across colony biofilm models in the symbiont *Vibrio fischeri*

**DOI:** 10.1101/2023.08.07.552283

**Authors:** Jacob A. Vander Griend, Ruth Y. Isenberg, Ketan R. Kotla, Mark J. Mandel

**Affiliations:** Department of Medical Microbiology and Immunology, University of Wisconsin-Madison, Madison, WI USA; Microbiology Doctoral Training Program, University of Wisconsin-Madison, Madison, WI USA

**Keywords:** bacterial biofilm, symbiosis, regulatory pathways, phosphorelay, two-component signaling systems, comparative transcriptomics

## Abstract

Beneficial microbial symbionts that are horizontally acquired by their animal hosts undergo a lifestyle transition from free-living in the environment to associated with host tissues. In the model symbiosis between the Hawaiian bobtail squid and its microbial symbiont *Vibrio fischeri,* one mechanism used to make this transition during host colonization is the formation of biofilm-like aggregates in host mucosa. Previous work identified factors that are sufficient to induce *V. fischeri* biofilm formation, yet much remains unknown regarding the breadth of target genes induced by these factors. Here, we probed two widely-used *in vitro* models of biofilm formation to identify novel regulatory pathways in the squid symbiont *V. fischeri* ES114. We discovered a shared set of 232 genes that demonstrated similar patterns in expression in both models. These genes comprise multiple exopolysaccharide loci that are upregulated and flagellar motility genes that are downregulated, with a consistent decrease in measured swimming motility.

Furthermore, we identified genes regulated downstream of the key sensor kinase RscS that are induced independent of the response regulator SypG. Our data suggest that putative response regulator VpsR plays a strong role in expression of at least a subset of these genes. Overall, this study adds to our understanding of the genes involved in *V. fischeri* biofilm regulation, while revealing new regulatory pathways branching from previously characterized signaling networks.

**IMPORTANCE:** The *V. fischeri-*squid system provides an opportunity to study biofilm development both in the animal host and in culture-based biofilm models that capture key aspects of *in vivo* signaling. In this work, we report the results of the transcriptomic profiling of two *V. fischeri* biofilm models followed by phenotypic validation and examination of novel signaling pathway architecture. Remarkable consistency between the models provides a strong basis for future studies using either–or both–approaches. A subset of the factors identified by the approaches were validated in the work, and the body of transcriptomic data provides a number of leads for future studies in culture and during animal colonization.

## INTRODUCTION

In animal-microbe mutualisms, symbionts can provide nutritional, developmental, and/or defensive benefits to the host, and in turn, the symbionts may gain various benefits from association with the host (1–6). During horizontal transmission, hosts must reacquire their symbionts each generation from environmental symbiont populations (7–14). Unfortunately, the understanding of how specific microbes make this transition from environment to host is often hindered by the complexity of animal microbiomes (15, 16). Symbioses with limited symbiont diversity are therefore valuable as models to identify mechanisms of colonization and transmission. The binary mutualism between the nocturnal Hawaiian bobtail squid (*Euprymna scolopes*) and the bioluminescent marine bacterium *Vibrio fischeri* is one such model system that integrates symbiont specificity, a defined colonization program, and a genetically tractable microbe (14, 17, 18).

Upon hatching from aposymbiotic (symbiont-free) eggs, juvenile squid rapidly begin symbiont recruitment from the surrounding seawater (18, 19). Located in the squid’s mantle cavity, the bilobed symbiotic “light organ” actively captures bacteria from seawater through the activity of extruded ciliated appendages on each lobe, which focus water currents onto a ciliated mucosal layer on the exterior of the organ (20). *V. fischeri* cells that become entrained in host mucosa form biofilm-like aggregates through the secretion of a specific exopolysaccharide (13, 21, 22). Aggregate formation is a critical step in host colonization, and mutants defective in aggregation are generally compromised in reaching the internal crypt spaces of the light organ where the symbiosis is maintained (22, 23). Therefore, understanding the genetic program connected to *in vivo* biofilm formation is critical to understand the transition that the colonizing microbes undergo from the planktonic state in seawater to successfully engraft themselves into the host microbiome.

*V. fischeri* aggregates have been shown to require production of the <u>sy</u>mbiosis exo<u>p</u>olysaccharide (Syp), produced and exported by the products of a conserved 18-gene (*syp*) locus (21, 22, 24, 25). Control of the *syp* locus is principally coordinated through a phosphorelay network feeding into two downstream response regulators (21, 23, 26) (**Fig. 1A**). Following auto-phosphorylation of the hybrid sensor kinase RscS, the phosphoryl group is transferred to the hybrid sensor kinase SypF (21, 27). Upon phosphorylation of SypF, downstream phosphotransfer events are targeted from SypF onto the response regulators SypE and SypG (27). SypG is a response regulator and ☐^54^-dependent activator that, when phosphorylated in its receiver (REC) domain, binds four promoters within the *syp* locus and activates their transcription (28). In contrast to the DNA-binding functionality of SypG, SypE is believed to act as a post-transcriptional regulator of Syp production that acts through SypA (29). As an additional level of control, *V. fischeri* also inhibits expression of the *syp* locus through the action of the biofilm inhibitor sensor kinase (BinK), which acts through SypG (30, 31).

Despite the robust induction of Syp exopolysaccharide production during host colonization, *in vitro* (i.e., culture-based) models of *V. fischeri* require genetic manipulation or chemical supplementation to induce biofilm formation (23, 32–34). One method to induce biofilm is via overexpression of the sensor kinase RscS, either through the plasmid-based *rscS1* allele or the chromosomal *rscS\** allele (21, 35, 36). RscS overexpression also induces increased aggregate formation in the squid host, explicitly linking the ability of this model to form biofilms *in vitro* with aggregation in the host context (21). Also, deletion of the gene encoding inhibitor sensor kinase BinK was also found to induce significantly larger aggregates during colonization (30). In culture, this biofilm-up phenotype can be reproduced by treating a Δ*binK* mutant with levels of calcium comparable to those found in seawater (Δ*binK-*Ca^2+^), which similarly results in wrinkled colony biofilm formation (33). Both of the culture-based *rscS\** and Δ*binK*-Ca^2+^ models increase biofilm formation by stimulating the expression of the *syp* locus through the sensor kinase SypF (27, 33). However, the models differ in the input by which SypF is phosphorylated, with RscS acting as the primary phosphodonor to SypF in the *rscS\** model, while the Δ*binK-*Ca^2+^ model requires a secondary phosphorelay involving the sensor kinase HahK (Δ*binK-*Ca^2+^) (27, 33).

**Figure 1.**
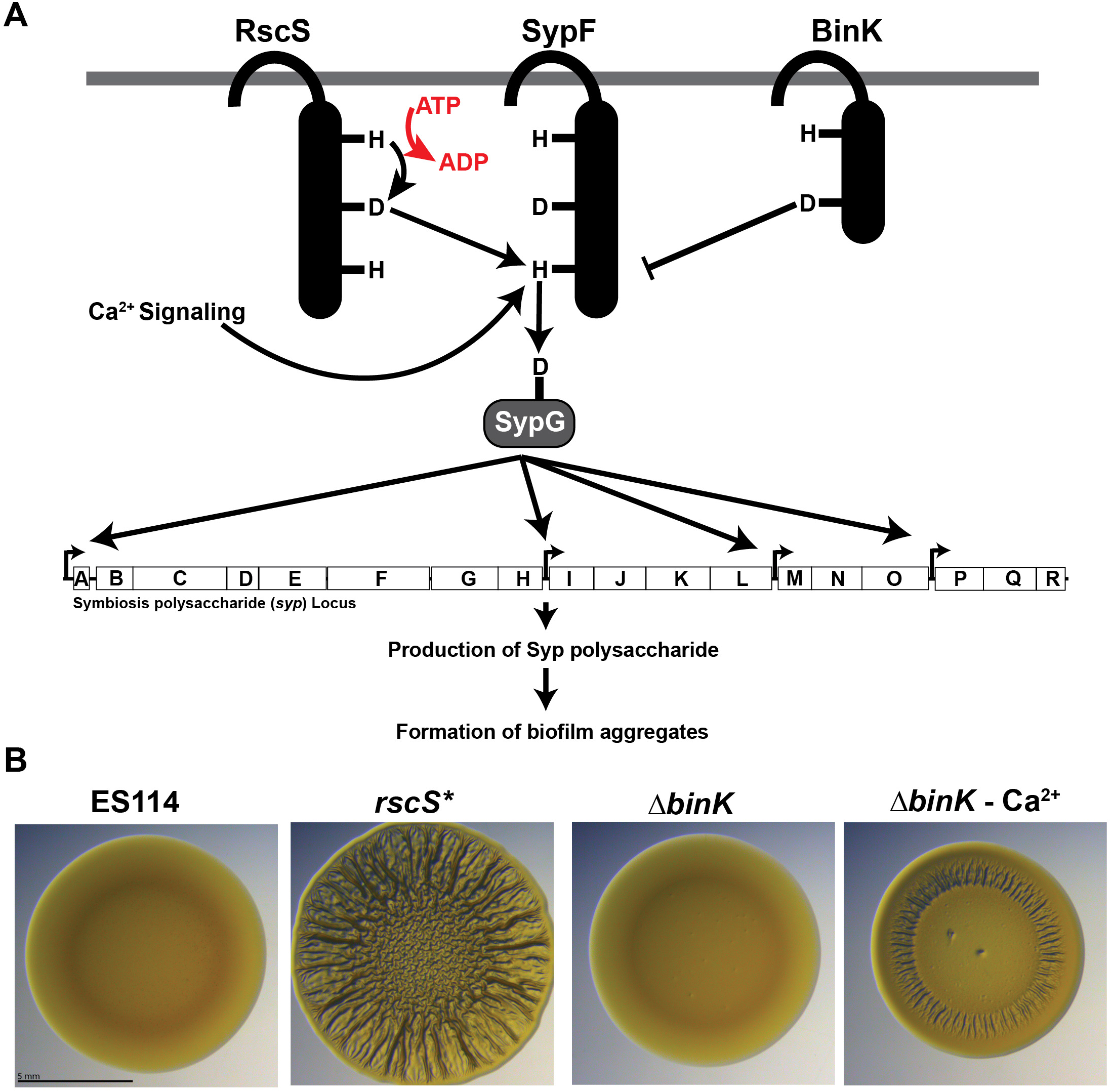
*Vibrio fischeri* symbiosis polysaccharide regulation. **A.** *Vibrio fischeri* induces biofilm formation through a phosphorelay initiated at the hybrid sensor kinase RscS, flowing through the hybrid sensor kinase SypF to the response regulator SypG. Growth on calcium media induces a secondary pathway also flowing through SypF. In opposition to this pathway, the sensor kinase BinK inhibits the activation of SypG, preventing biofilm formation. **B.** *V fischeri* ES114 has a smooth colony morphology when grown on LBS-agar. Overexpression of the sensor kinase RscS *(rscS*)* or growth of a *t,.binK* mutant on LBS 10 mM Ca^2+^ agar permit the formation of "wrinkled" colonies with a rugose morphology, due to production of the Syp exopolysaccharide.

The discovery of both the RscS overexpression and Δ*binK*-Ca^2+^ models opened the door to genetic analysis of Syp regulation, as both models recapitulate the induction of the critical Syp exopolysaccharide without requiring the squid host (27, 30, 33, 37). Induction of the Syp exopolysaccharide in culture manifests as a distinct wrinkling morphology within colonies grown on agar media, while the wild type strain of *V. fischeri* ES114 remains smooth (**Fig. 1B**) (24, 33). Considering the shared dependence of both wrinkled colony biofilm models and *in vivo* aggregates on Syp exopolysaccharide production, in this work we utilized wrinkled colonies formed by both of the biofilm models as proxies for *in vivo* aggregates in a comparative transcriptomics assay.

## RESULTS

### Definition of a core biofilm regulon for *V. fischeri* ES114

*V. fischeri* wrinkled colony biofilms are a close proxy for the aggregates that form *in vivo* during squid host colonization (21, 22). To begin to understand the genes that are induced under these conditions, we asked what patterns of gene expression were shared in two common wrinkled colony biofilm models: (i) Genetic induction with an *rscS* overexpression allele (*rscS\**) in an otherwise wild type background compared to wild-type *V. fischeri* ES114, and (ii) Chemical induction with 10 mM CaCl_2_ in a Δ*binK* background (Δ*binK*-Ca^2+^) compared to Δ*binK* alone (33, 36) (**Fig. 1B**). We selected these comparisons to explicitly compare biofilm induced to the closest uninduced control conditions for each. To query gene expression in wrinkled colony biofilms, we compared the global transcriptome in the induced condition with the uninduced condition in each case. Spots of overnight culture were spotted onto fresh agar and incubated at 25 °C to form wrinkled colony biofilms. After 48 h, each spot was scraped, and RNA was isolated for differential expression analysis. Throughout the study, we used thresholds of 2-fold induction/repression and false discovery rate (FDR) of 0.05.

***(i) Effects on exopolysaccharide gene transcription*** In both biofilm models, we detected a large number of genes that were differentially expressed, with over 200 genes upregulated and over 200 genes downregulated upon biofilm induction (**Fig. 2A**). Comparing the results from the two models revealed 123 genes that were significantly upregulated and 109 genes that were significantly downregulated in both cases (**Fig. 2B)**. The most highly-upregulated genes included those in the symbiosis exopolysaccharide (*syp*) locus, which are required for aggregation in the host and are known to be regulated downstream of RscS and Δ*binK*-Ca^2+^ signaling (33) (**Fig. 2C)**. Examination of the data revealed a remarkably consistent response for the *syp* genes across both models. The *syp* locus is organized into four operons, each of which has a σ^54^ promoter and binding sites for the enhancer-binding protein SypG to facilitate transcription by Eσ^54^ (28). We noted a trend in which the first gene in each operon was the most highly-induced, with *sypA* being the highest induced gene across the entire locus with over 97-fold induction in the *rscS\** background and 49-fold induction in the Δ*binK*-Ca^2+^ background (**Fig. 2D**). Again, responses under the two biofilm models were highly correlated. The three SypG-regulated biofilm maturation (matrix) proteins (BMP) and three biofilm-associated lipoproteins (BAL) are similarly regulated by both models (**Fig. 2C**) (38). These data therefore illustrate that our RNA-seq analysis from agar-grown colony spots capture relevant transcriptional responses and that integration of both models yields valuable signals. In the remainder of the study, we refer to genes that show consistent regulation under both models as the “core biofilm regulon”. One unexpected result was that *sypG* had the lowest biofilm induction among the 18 *syp* genes. Given the importance of SypG to biofilm regulation as the σ^54^-dependent activator for transcription of the *syp* promoters, we were curious to investigate the relative transcript levels of *sypG* within a sample, especially in the uninduced conditions. We conducted an analysis of gene expression within each condition using transcripts per kilobase million reads (TPM) and focused on the genes within the *syp* locus (39). This analysis revealed that *sypF* and *sypG* had the highest TPM values in the uninduced wild-type sample (**Fig. S1**). When we similarly examined the induction of these genes in the *rscS\** and Δ*binK*-Ca^2+^ models, we noted similar patterns of expression, with SypF and SypG appearing to break the decreasing pattern of expression across operon 1. These results argue that *sypF* and *sypG* have higher basal transcriptional levels in non-biofilm induced conditions and suggest that they may be regulated in a distinct manner from the other genes across the locus. Beyond the *syp* locus, *V. fischeri* encodes two additional exopolysaccharide loci (26, 40). The products of the *bcs* locus produce cellulose, which permits adhesion to surfaces and may regulate the transcription of the *syp* locus (26, 41). In both biofilm models we observed moderate upregulation of *bcs* genes (**Fig. S2A**), with *bcsQ* (*VF_A0885*) and *bcsE* (*VF_A0886*) exhibiting the highest fold changes, fitting the *syp* pattern of the first gene in each respective operon being the most highly upregulated. The other exopolysaccharide locus spans *VF_0157-VF_0180* and is regulated by the quorum sensing regulator LitR (40). To date, the function of this locus remains unclear, yet it does appear to enhance phage infection by the *V. fischeri* phage HNL01 (6). In contrast to the upregulation of the *bcs* and *syp* loci within the core biofilm regulon, we noted that a majority of genes in the *VF_0157-VF_0180* EPS locus were downregulated modestly in both models (**Fig. S2B**).
***(ii) Effects on other biofilm-related loci of note***. Among the most-highly upregulated genes in the core biofilm regulon, we noted a predicted curli amyloid fiber biosynthetic locus (**Fig. 3A**). In other bacteria, CsgA subunits polymerize into curli amyloid fibers on the exterior of the cell in a process nucleated by CsgB (42, 43). Curli fibers mediate aggregation and biofilm formation in different species, and the operon structure in *V. fischeri* mimics that of other organisms including predicted export genes (43–45). The *V. fischeri* CsgA and CsgB proteins are larger than their *Escherichia coli* homologs, and each share 32.0 % similarity, respectively, to the characterized curli proteins from *E. coli* K-12 (46, 47). Notably, *V. fischeri* curli proteins are larger than their *E. coli* homologs, with CsgA and CsgB predicted to be 315 amino acids and 190 amino acids, respectively. We generated a strain lacking the *csgBA* genes (Δ*csgBA*), eliminating both the major and minor curlin subunits, and queried whether interruption of the curli genes reduced host colonization levels. We observed no significant difference in final colonization levels between wild type and the Δ*csgBA* mutant (**Fig. 3B**), suggesting that the curli locus is not a critical colonization factor in this context. We further tested whether the absence of curli would impact biofilm aggregate formation in the host. However, we did not observe any significant reductions in the size of biofilm aggregates in the *ΔcsgBA* mutant compared to wild type, suggesting that the curli locus is not required in host biofilm aggregates (**Fig. 3C**). Outside of the squid host, we tested whether the curli amyloid fiber locus impacted *in vitro* models of biofilm formation by introducing the *ΔcsgBA* mutation into a RscS overexpression background (*rscS\**). We noted no significant defects in wrinkled colony formation in the *rscS\** Δ*csgBA* mutant at both early (24 h) and late (48 h) timepoints (**Fig. 3C**). Additionally, the Δ*csgBA* mutant did not have decreased binding of the amyloid-binding dye Congo Red, which is notably different to deletions of *csgBA* genes in other organisms (48). Given the linkage between curli production and cellulose EPS in other organisms (49, 50), we proceeded to test whether deletion of both *csgBA* and the cellulose EPS synthase *bcsA* would impact biofilm formation. In the wrinkled colony biofilm model, the strain lacking both curli and *bcsA* demonstrated no significant differences in morphology compared to a parental biofilm induced strain, and it exhibited Congo Red binding equivalent to a *bcsA* single mutant. Therefore, while it is intriguing that the curli system is induced concomitant with biofilm formation, we did not detect a functional requirement for the curli genes. Despite the similarity in operon structure, we do note that the *V. fischeri* CsgA and CsgB proteins are only 32% similar and substantially larger than their *E. coli* K-12 orthologs (2.1x and 1.25x in length, respectively) (46, 47). Given our results and the lack of Congo Red binding attributable to the curli gene products, we are unable to determine whether curli fibers are produced or play a role during symbiotic colonization.
***(iii) Effects on flagellar motility***. Outside of exopolysaccharide regulation, the core biofilm regulon had a substantial downregulation of motility genes. We observed that 32 of the 43 genes required for swimming motility in *V. fischeri* (51) trended towards reduced expression in both biofilm models. Of these genes, five met our significance threshold: *cheB/VF_1830*, *cheY/VF_1833*, *fliN/VF_1844*, *fliM*/*VF_1845*, and *flrB/VF_1855* (**Fig. 4A**, **Table S1**). We therefore asked whether the *rscS\** or Δ*binK* backgrounds had a deficit in swimming motility by examining migration in TBS soft agar swim plates (**Fig. 4B**). Compared to wild type controls, either *rscS\** or the Δ*binK* genetic backgrounds alone were sufficient to reduce swimming migration on soft agar by at least 15%, demonstrating that upregulation of the biofilm pathway leads to a diminution of swimming motility (**Fig. 4C**).
***(iv) Effects on cyclic-di-GMP***. In *V. fischeri* and in many bacteria, increased cyclic-di-GMP (c-di-GMP) levels positively impact the production of the cellulose exopolysaccharide and negatively regulate swimming motility, both effects noted in our study (41, 52). Within the transcriptomics dataset, we asked which diguanylate cyclases (DGCs) and phosphodiesterases (PDEs) were differentially regulated under these conditions (**Fig. S3A, Table S2**). Both biofilm backgrounds upregulated two DGCs (*VF_A0152* and *casA/VF_1639*), two PDEs (*VF_A0526* and *binA*/*VF_A1038*), and one predicted bifunctional DGC/PDE (*VF_0985*). In contrast, both backgrounds downregulated four DGCs (*VF_1350*, *VF_A0398*, *mifB*/*VF_A0959*, and *VF_A1012*), one PDE (*VF_A0551*) and two DGC/PDEs (*VF_0094* and *VF_A0244*). We next directly tested production of c-di-GMP using a plasmid-based fluorescent reporter (pFY4535) (53). This reporter has constitutive expression of the AmCyan fluorescent protein, with the expression of TurboRFP under the control of c-di-GMP binding riboswitches. Higher levels of c-di-GMP therefore increase the TurboRFP/AmCyan ratio. Compared to wild type *V. fischeri* and control strains known for either high c-di-GMP levels (Δ6PDE) or low cyclic-di-GMP levels (Δ7DGC) (41, 52), we noted that *rscS\** induced a mild but significant increase in c-di-GMP levels in colony biofilms grown on standard LBS media, while no such induction was observed in the Δ*binK* background (**Fig. S3B**). Calcium supplementation (10 mM) of LBS media greatly increased cyclic-di-GMP levels in both *rscS\** and the Δ*binK* colony biofilms to equivalent levels. This result may be explained by the shared upregulation of the calcium-induced DGC *casA* in both models, as overexpression of CasA has previously been linked to increased c-di-GMP synthesis in calcium supplemented liquid cultures (**Fig. S3C**) (54). Overall, both models of biofilm formation appear to be capable of increasing the global levels of c-di-GMP.

**Figure 2.**
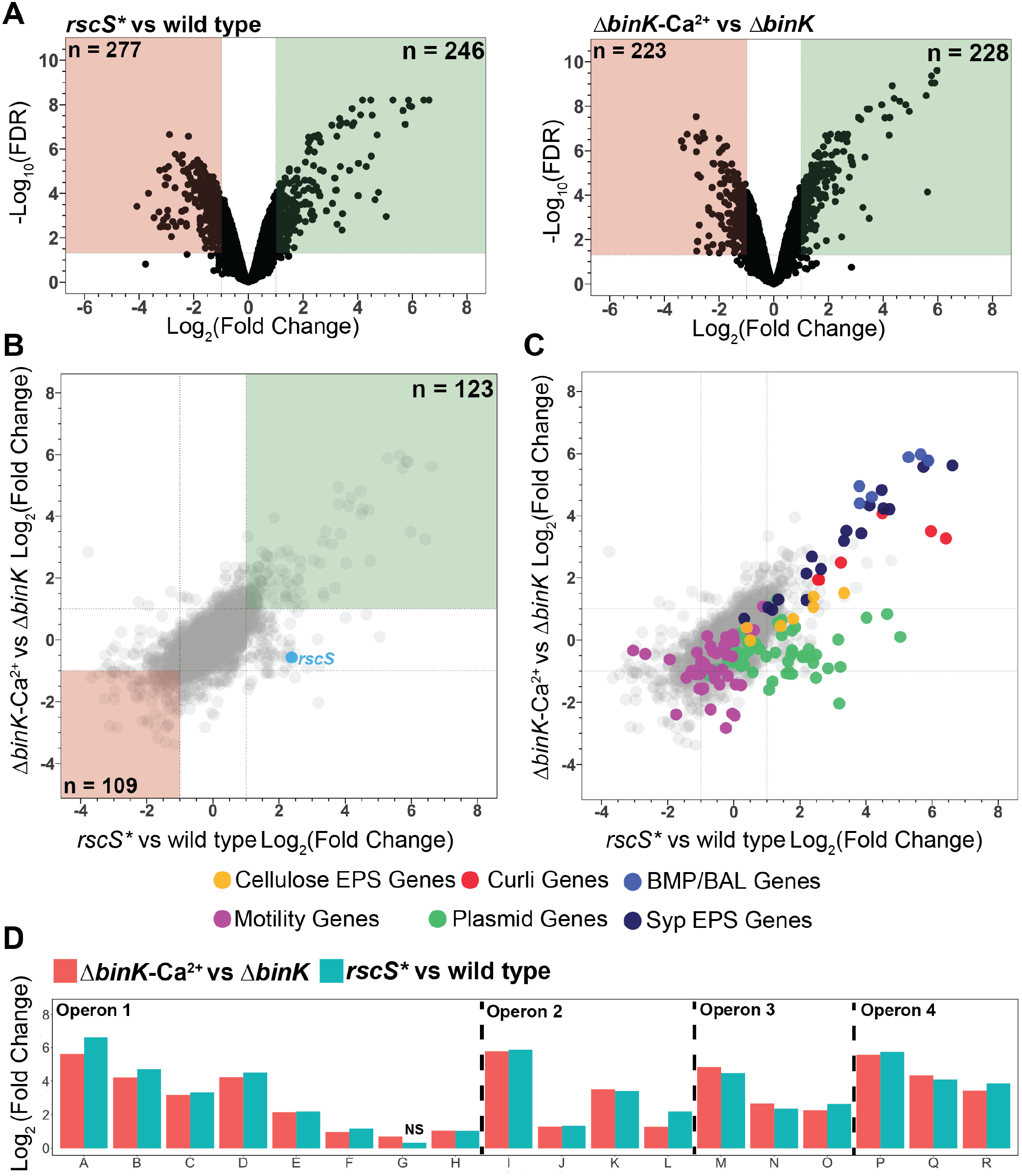
*Vibrio fischeri* induces a core transcriptomic response during biofilm formation. **A.** Volcano plots of the *rscS\** vs wild type and *!).binK-Ca*^2^*^+^* vs *!).binK* biofilm models. Green and red boxes indicate those genes with LogiFold Change) > 1 and < -1 respectively, with significant differential expression (FDR< 0.05). **B.** Overlay of LogiFold Change) values for all genes from both the *rscS\** vs wild type and *!).binK-Ca*^2^*^+^* vs *!).binK* biofilm models. Green and red boxes are analogous to those in A. **C.** Overlay of annotations onto the multi-comparison overlay shown in B, dot legend is provided below. **D.** Syp locus differential expression. LogiFold Change) for each of the 18 genes in the *syp* locus is provided from the two differential expression analyses. Operon indications are provided from previous analyses of SypG-bound promoters (28). For the one gene which had no significant changes in expression *(sypG; rscS\** vs wild type), NS is indicated above that column.

**Figure 3.**
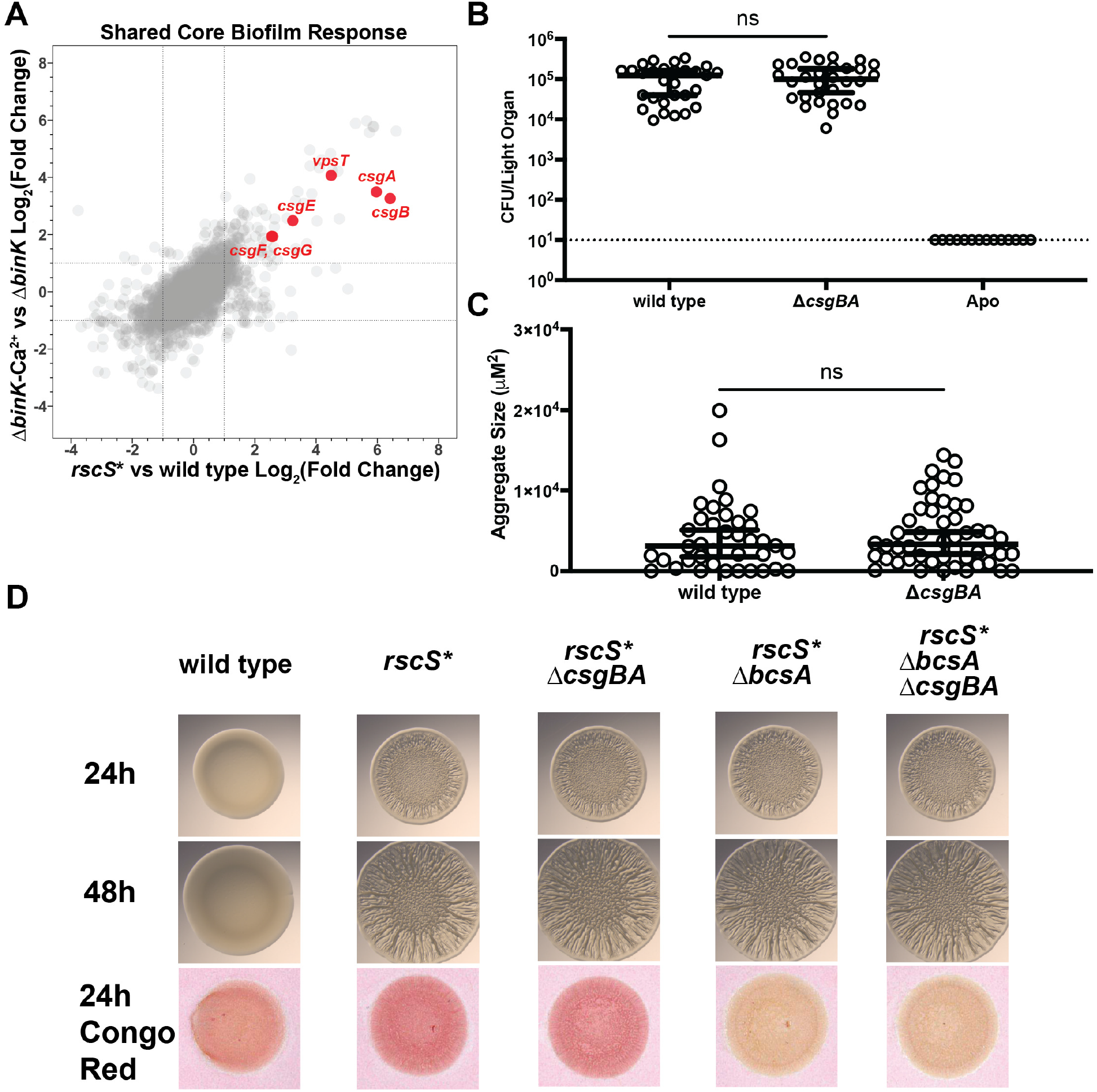
Curli locus is highly induced by both biofilm models. **A.** Overlay graph of Fig. 2C with specific curli genes (red), labeled by name. **B.** Single strain colonization data for wild type and a *ticsgBA* curli mutant. Wild type and *ticsgBA* were inoculated into filter sterilized Instant Ocean (FSIO) of two separate cohorts of squid hatchlings, and allowed to colonize for 3 h. Hatchlings were then washed and allowed to grow for 2 d, after which squid juveniles were euthanized. Each dot represents the average CFU/ml of individual squid, and horizontal bars represent median CFU/ml with 95% confidence intervals. The dashed line indicates the limit of detection. Statistical significance was calculated using a Mann-Whitney test. (ns, not significant). **C.** Biofilm aggregate size during host colonization. Wild type and *ticsgBA* strains carrying the constitutive GFP plasmid pVSV102 were allowed to colonize squid hatchlings for 3 hours, at which point each squid was euthanized and fixed in 4% paraformaldehyde. Dissection of each squid then revealed the biofilm aggregates forming on the light organ surface, which were imaged with a Zeiss Axio Zoom fluorescence microscope. Each point represents the area of one aggregate. Squid which did not have any aggregates have points at zero. Horizontal bars represent median aggregate area size with 95% Cl. Statistical significance was calculated using a Mann-Whitney test. (ns, not significant). **D.** Wrinkled colony and Congo red binding assays. Strains were spotted on LBS media and imaged.

**Figure 4.**
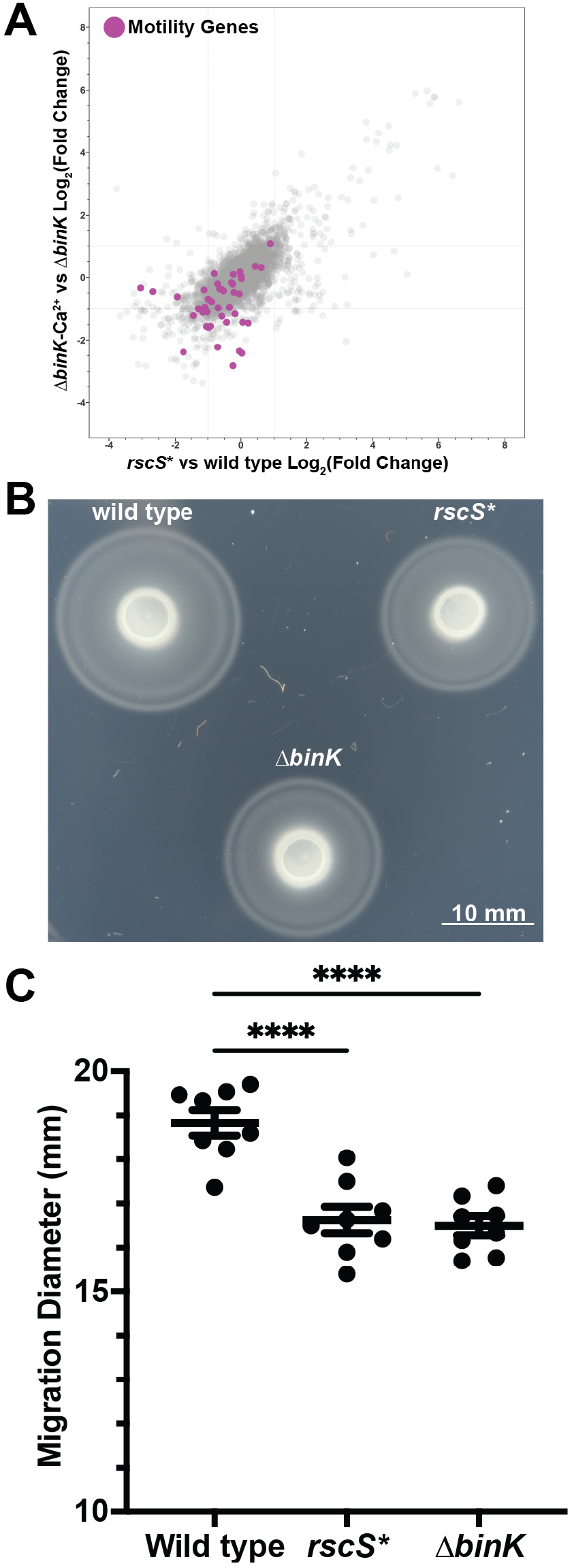
Biofilm-induced conditions downregulate flagellar motility genes and have reduced motility. **A.** Both biofilm-induced models significantly downregulate genes required for flagellar biosynthesis genes from Brennan *et al*. (51). **B.** Strains were cultured in TBS media, spotted onto TBS soft-agar plates, and incubated at 25 °C for 5 h. Shown is a representative plate of n = 8 biological replicates, imaged with a Nikon D810 camera. **C.** Migration distance determined using lmageJ. Statistical significance was calculated using an ordinary one-way ANOVA using Dunnett’s multiple comparisons test in GraphPad Prism. Each dot is the product of n = 3 technical replicates, horizontal lines represent the mean with SEM (****, P **<** 0.0001).

### RscS overexpression and calcium supplementation induce regulatory programs unique to each model

Despite the contributions of both biofilm models to the shared biofilm regulon, we also noted targets that appeared to be regulated by either model alone. For the RscS overexpression model, we detected 61 genes with increased expression and 68 genes with reduced expression (**Table S3**). Notable targets within the upregulated group included genes encoding the regulator HbtR/*VF_A0473* and its chaperone HbtC/*VF_A0474*, alongside 15 genes encoded on the natural plasmid of *V. fischeri* ES114, pES100. Of the 68 downregulated genes, we observed multiple genes associated with flagellar motility including *fliA/VF_1834*, flagellar motor proteins *motA2/VF_A0186* and *motB2/VF_A0187*, and the chemotaxis factors *cheV2/VF_A0802* and *VF_2042*. Compared to the RscS overexpression model, the Δ*binK*-Ca^2+^ model affected fewer targets, with 38 upregulated genes and 53 downregulated genes specific to this model (**Table S3**). The upregulated group contained multiple iron and copper assimilation loci and the quorum sensing autoinducer C8-HSL synthase AinS. Downregulated genes in this model included multiple flagellar genes including *flaA/*VF_1866, as well as predicted regulatory factors including the cold-shock DNA-binding transcriptional regulator *cspG/*VF_A1094.

### Novel members of the core biofilm regulon are induced independently of the response regulator SypG

The dominant mode of signaling from sensor kinase RscS is through hybrid sensor kinase SypF to response regulator SypG (27). We therefore expected that removal of SypG in the *rscS\** overexpression background would eliminate most of the effect observed in that model. While the *syp, bmp*, and *bal* genes did not exhibit induction in the absence of SypG, many of the remaining *rscS\**-induced genes continued to be differentially expressed without the downstream regulator (**Fig. 5A**). 111 genes were significantly upregulated and 60 genes were significantly downregulated, despite the attenuation of the *rscS\** Δ*sypG* mutant in forming wrinkled colony biofilms. These results suggest that RscS maintained a regulatory program independent of SypG signaling (**Fig. 5B**). Visualization of these data reveal a clear bifurcation showing this unexpected SypG-independent effect within the *rscS\** model (**Fig. 5C, Table S4**).

**Figure 5.**
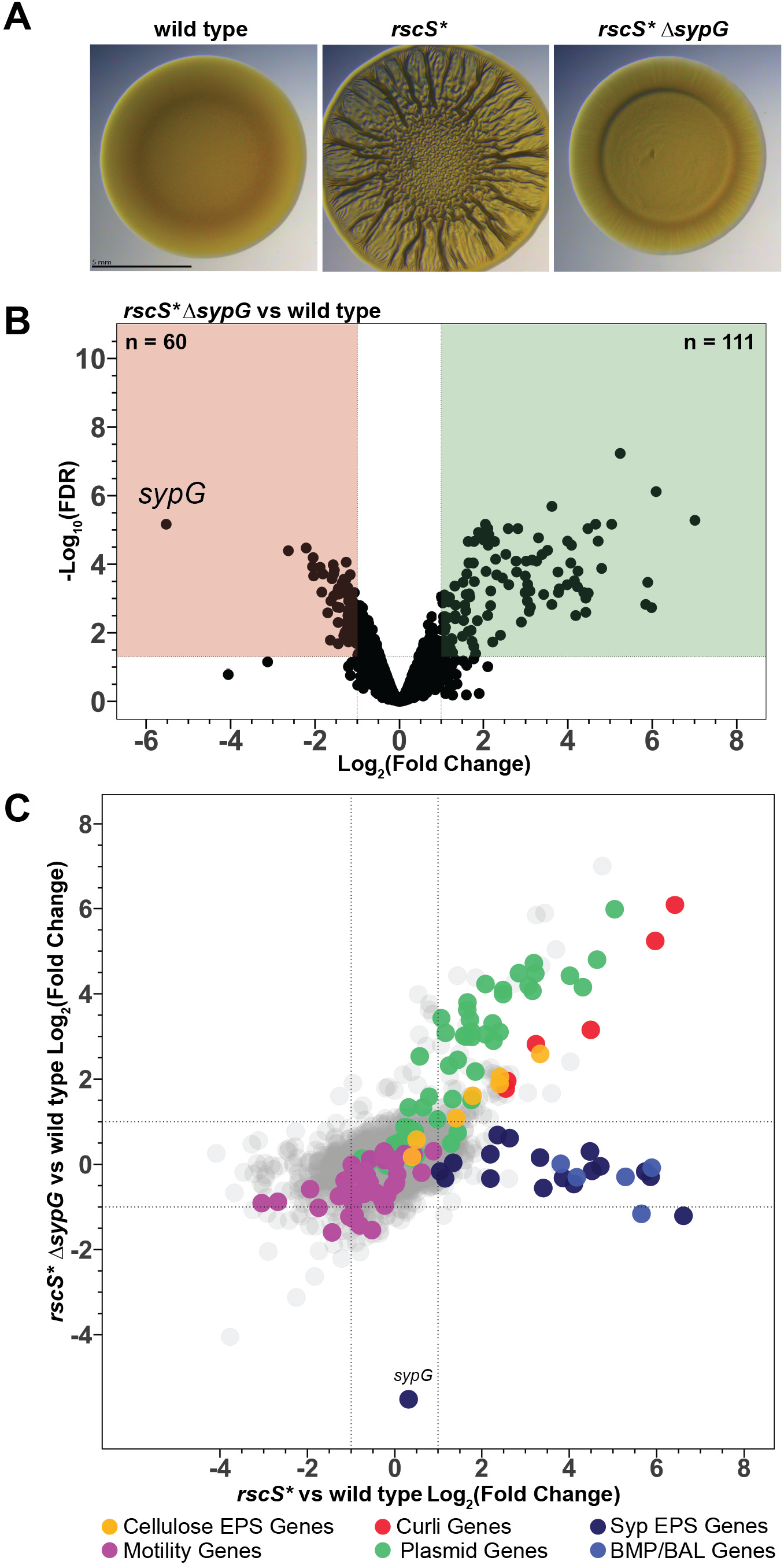
Deletion of the response regulator gene *sypG* reveals a signaling bifurcation downstream of RscS. **A.** Wrinkled colony assay. **B.** Volcano plot of wild type versus *rscS* 1’1.sypG* differential expression analysis. Green and red boxes demarcate genes with Log/Fold Change) ;;:: 1 and::; -1 respectively, with significant differential expression (FDR< 0.05). The number of genes in each box are listed on the figure. **C.** Overlay of Log/Fold Change) values for all genes in both the *rscS\** vs wild type and the *rscS* 1’1.sypG* vs wild type models.

The SypG-independent signaling pathway regulates *bcs* locus transcription through the response regulator VpsR and independent of SypF. The detection of SypG-independent RscS regulation suggested that RscS was not limited to being simply a regulator of the *syp* locus. However, the factors involved in transmitting RscS signaling to alternative pathways remained unclear. One possibility was that RscS might branch a new signaling pathway through its partner sensor kinase SypF. SypF has precedent as a sensor kinase with multiple downstream response regulators, affecting both SypE and SypG within the Syp phosphorelay (27, 29). Additionally, an increased activity allele of SypF (SypF1) was discovered to increase production of the cellulose exopolysaccharide (26, 54). This activity required the gene *VF_0454*, which encodes a homolog of the *V. cholerae* response regulator VpsR (26, 54). As we observed robust induction of the cellulose exopolysaccharide (*bcs*) locus in the *rscS\** Δ*sypG* background (versus wild type) and in the core biofilm regulon, we hypothesized that SypF-mediated VpsR signaling could explain the SypG-independent signaling pathway. To test this hypothesis, we asked whether transcription of SypG-independent gene *bcsE* was SypF- and VpsR-dependent. Using fluorescence microscopy, we measured reporter expression in wrinkled colony biofilms grown on LBS or LBS-Ca^2+^ agar. While removal of SypF or SypG exhibited mild effects (up to 18%), only removal of VpsR eliminated the induction of the reporter (**Fig. 6AB**). Testing another SypG-independent promoter, for *VF_0208*, revealed roles for SypF, SypG, and VpsR, yet again VpsR exhibited the strongest impact on the promoter’s activity (**Fig. 6CD**). Together, these data suggest that VpsR plays an important role in the regulation of genes in the *rscS\** model.

**Figure 6.**
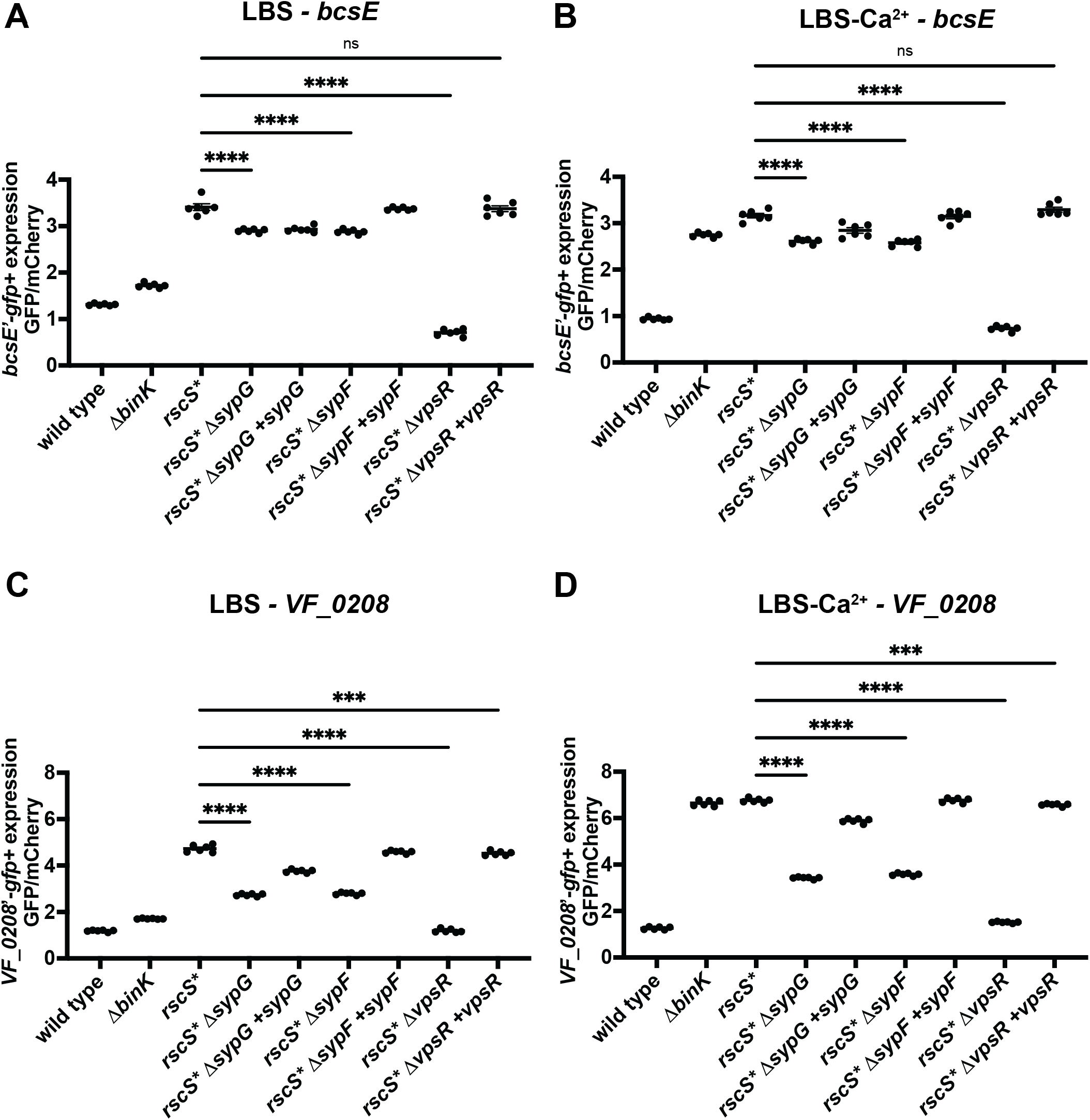
RscS uses VpsR to regulate SypG-independent targets. **A.** Expression data for the *bcsE’-gfp^+^* reporter for strains spotted on LBS media. Complementation backgrounds are indicated by the presence of *+vpsR, +sypF,* and *+sypG,* all complementations were single copy insertions at the *Vfischeri* Tn7 transposon site. Fluorescence was measured within colony spots with *n* = 3 biological replicates, each the product of n = 2 technical replicates using a Zeiss Axio Zoom fluorescence microscope. GFP fluorescence was normalized to mCherry fluorescence to determine the expression of the reporter in each strain. Each dot represents one biological replicate, each the product of n = 2 technical replicates, collected from experiments on two separate days. Statistical analysis of each strain’s reporter expression was conducted in GraphPad Prism, using an ordinary one-way ANOVA, with Dunnett’s multiple comparison test. (ns, not significant; ***, *P* :5 0.001; ****, *P* < 0.0001) **B.** Expresion data for the *bcsE’-gfp^+^* reporter for strains spotted on LBS-Ca^2+^ media, collected and analyzed as in A. **C.** Expression data for the *vf_0208’-gfp^+^* reporter for strains spotted on LBS media, collected and analyzed as in A. **D.** Expression data for the *vf_0208’-gfp^+^* reporter for strains spotted on LBS-Ca^2+^ media, collected and analyzed as in A.

## DISCUSSION

In this work, we profiled two separate models of biofilm formation in the model symbiont *V. fischeri* ES114. This analysis revealed remarkable similarities between transcriptional responses to the two models, which facilitated identification of a core transcriptional program. In addition to known factors that were among the highest expressed in both models (e.g., *syp* locus genes), we identified novel regulatory targets that included well-annotated genes (e.g., motility), loosely-annotated genes (e.g., *csg, VF_0157-VF_0180* EPS), as well as many hypothetical and unannotated genes. This work generated testable hypotheses, and we pursued multiple examples of those within the study. We validated that in both models, biofilm formation resulted in diminished swimming motility, consistent with behavior in other organisms in which biofilm and motility are discordantly regulated (55, 56).

Our study provided insights into the quantitative nature of induction dynamics across biofilm operons. We observed the *syp* locus had dramatically higher induction for genes located earlier in each of the four operons, as seen for *sypA, sypI, sypM,* and *sypP* (**Fig. 2D**). This result suggests that *sypA* provides especially high dynamic range to use for reporting on core biofilm responses in strain ES114, consistent with previous studies that have applied *sypA’-gfp^+^* and *sypA’-lacZ^+^*transcriptional reporters (27, 31, 34, 35, 41). In uninduced conditions, we found that the genes encoding the central regulators of the Syp phosphorelay, *sypF* and *sypG,* had the highest abundance as measured by TPM among the *syp* gene transcripts. This result suggested to us that these transcripts may be regulated in a fashion uncoupled from the preceding genes. We speculate that there is a separate regulatory mechanism that enables a higher baseline of *sypF* and *sypG* transcripts without full *syp* locus induction, and that allows levels of SypF and SypG to accumulate to be able to respond to physiological induction of the pathway. In a recent study, induction of *sypF-H* in response to the biofilm stimulatory vitamin para-aminobenzoic acid (pABA) was observed in a pattern distinct from the rest of the locus (32), supporting a separate regulation mechanism that begins at *sypF*.

The large number of genes induced in the *rscS\** Δ*sypG* strain was surprising given that the strain does not induce biofilm formation in culture. This result suggested to us that there is a substantial novel output to the RscS signaling pathway, and in the two promoters we examined, we found that VpsR was required for activity in both cases. Mutants in *vpsR* exhibit a competitive defect in squid colonization and a defect in cellulose regulation in culture (26, 57). Despite characterized VpsR regulation via SypF, in the *rscS\** model we observed a substantial role for VpsR even when there was only a modest role for SypF (**Fig. 6AB**). We note that additional work on VpsR has been conducted in *V. cholerae*, which has a protein that is 66% identical to the *V. fischeri* ortholog. There, it has been shown to be regulated by phosphate and not by phosphorylation as would be expected as a putative response regulator (58).

Furthermore, in *V. cholerae* VpsR regulates the Vibrio polysaccharide locus (*vps*), which is absent in *V. fischeri*, as similarly the *bcs* locus is absent in *V. cholerae* (26). Overall, therefore, the role of VpsR in *V. fischeri* requires further clarification.

Overall, this work combines two models to identify novel aspects of biofilm regulation, reveal patterns of gene expression across regulated loci, and uncover new factors that are coregulated with the *V. fischeri* symbiotic biofilm program.

## MATERIALS AND METHODS

### Bacterial strains, plasmids, and media

*V. fischeri* and *E. coli* strains studied in this work are listed in **Table 1**. Plasmids generated or used in this work can be found in **Table 2**. Unless otherwise specified, *V. fischeri* strains were grown at 25 °C in Luria-Bertani salt (LBS) medium (per L: 25 g of Difco Miller LB broth [BD], 10 g of NaCl, 50 mL of 1M Tris buffer [pH 7.5], 10 mM CaCl_2_ [when noted]) or Tryptone Salt (TBS) medium (per L: 10 g of Gibco Bacto Tryptone, 20 g NaCl, 50 mL 1 M Tris buffer, pH 7.5, 3 g Agar). *E. coli* strains used for cloning and conjugation were grown at 37°C in Luria-Bertani medium (per L: 25 g of Difco Miller LB broth [BD]). When needed, antibiotics were supplemented at the following concentrations: Kanamycin (Kan), 100 µg/mL for *V. fischeri* and 50 µg/mL for *E. coli,* Chloramphenicol (Cam), 5 µg/mL for *V. fischeri* and 25 µg/mL for *E. coli*, Erythromycin (Erm) 5 µg/mL for *V. fischeri,* Gentamicin (Gent) 2.5 µg/mL for *V. fischeri* and 5 µg/mL for *E. coli.* When needed for specific strains, thymidine was added at 0.3 mM for *E. coli.* Solidified media was prepared with an agar concentration of 1.5% unless specified otherwise. For Congo red agar, 40 µg/mL Congo red and 15 µg/mL Coomassie blue were added to LBS. Plasmids were conjugated from *E. coli* strains into *V. fischeri* using standard techniques (59).

**Table 1.**
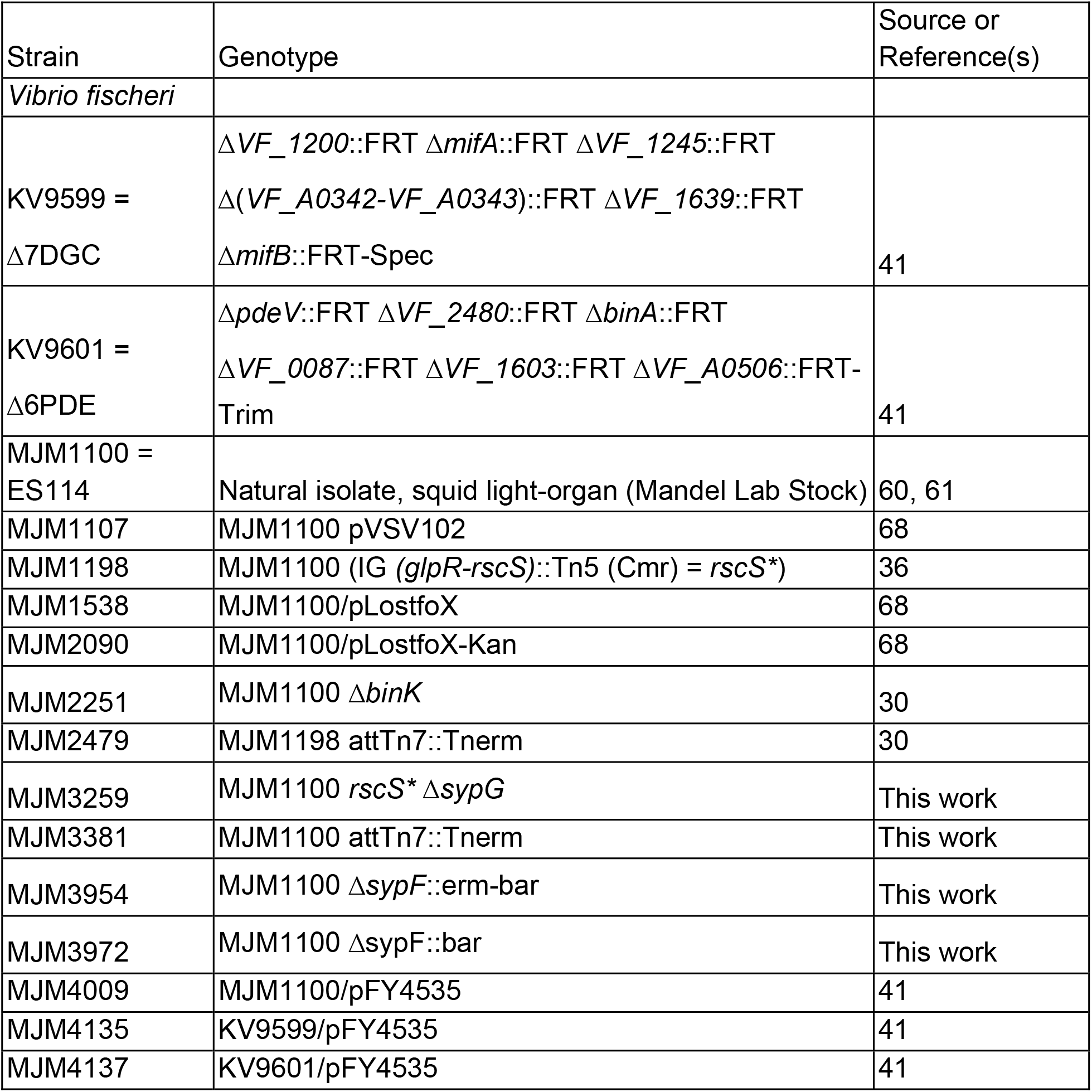
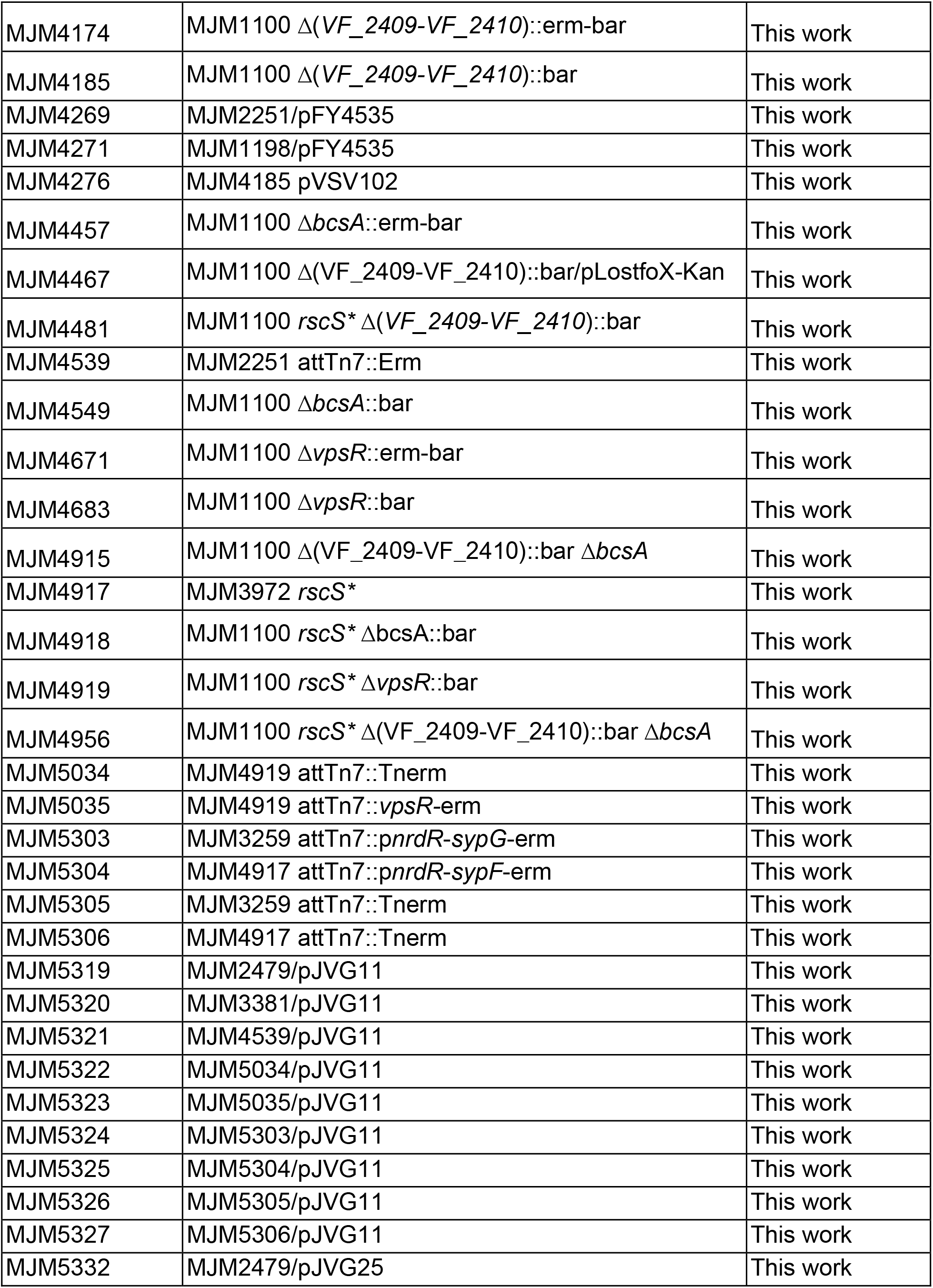
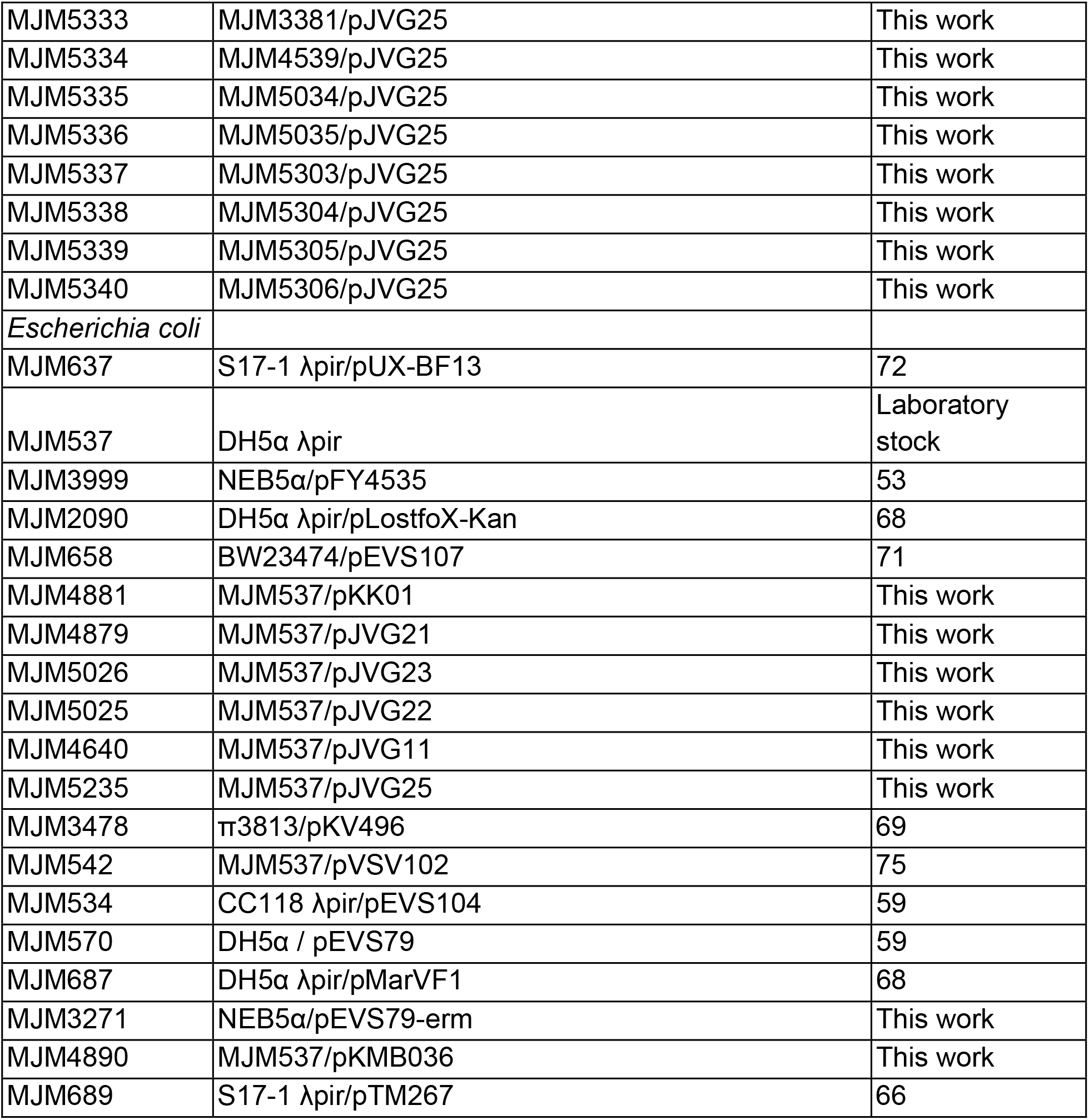
Strains:

**Table 2.**
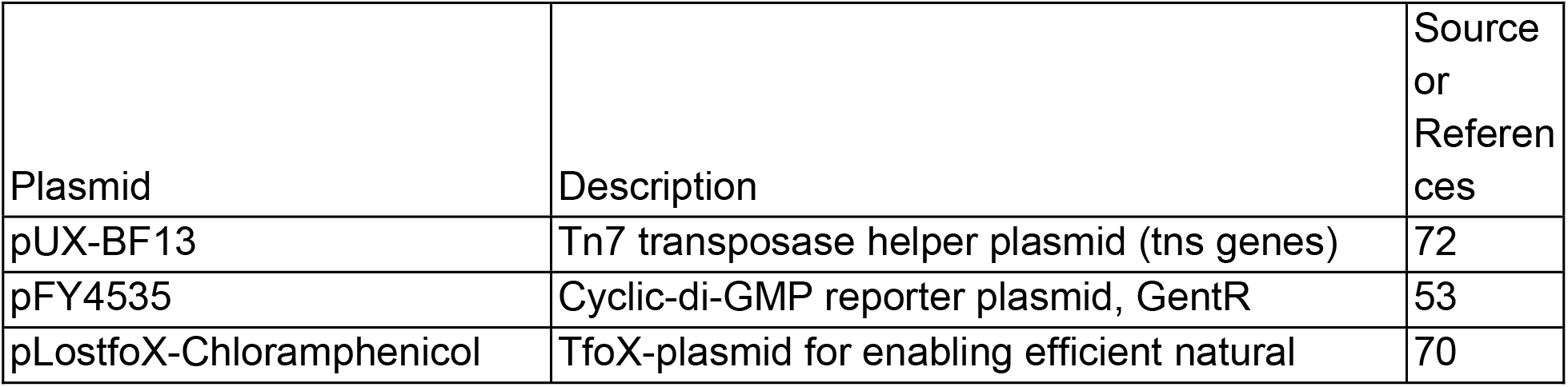
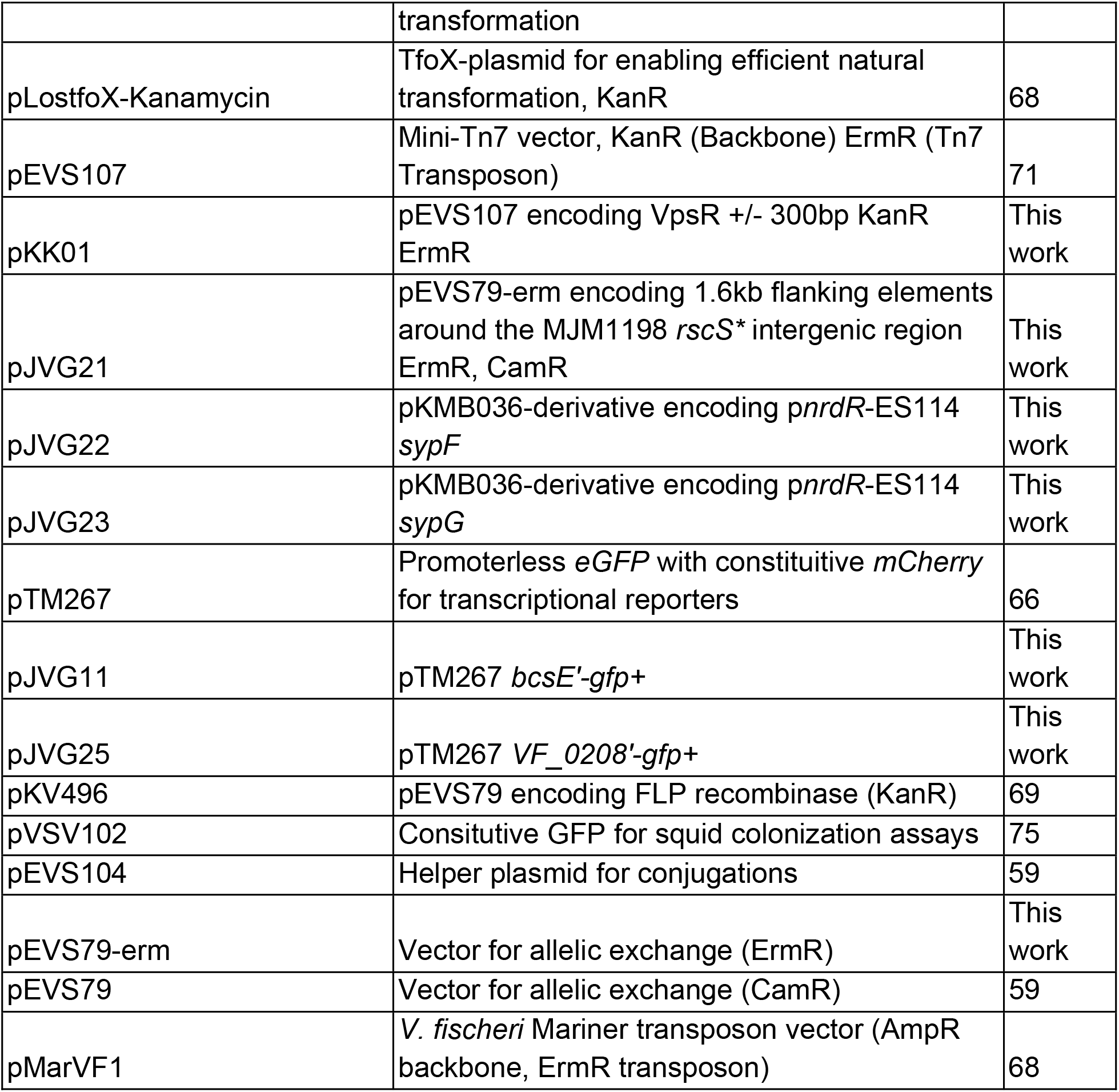
Plasmids:

| Plasmid | Description | Source or References |
| --- | --- | --- |
| pUX-BF13 | Tn7 transposase helper plasmid (tns genes) | 72 |
| pFY4535 | Cyclic-di-GMP reporter plasmid, GentR | 53 |
| pLostfoX-Chloramphenicol | TfoX-plasmid for enabling efficient natural | 70 |
|  | transformation |  |
| pLostfoX-Kanamycin | TfoX-plasmid for enabling efficient natural transformation, KanR | 68 |
| pEVS107 | Mini-Tn7 vector, KanR (Backbone) ErmR (Tn7 Transposon) | 71 |
| pKK01 | pEVS107 encoding VpsR +/- 300bp KanR ErmR | This work |
| pJVG21 | pEVS79-erm encoding 1.6kb flanking elements around the MJM1198 *rscS* * intergenic region ErmR, CamR | This work |
| pJVG22 | pKMB036-derivative encoding *pnrdr*-ES114 *sypF* | This work |
| pJVG23 | pKMB036-derivative encoding *pnrdr*-ES114 *sypG* | This work |
| pTM267 | Promoterless *eGFP* with constitutive *mCherry* for transcriptional reporters | 66 |
| pJVG11 | pTM267 *bcsE*'-gfp+ | This work |
| pJVG25 | pTM267 *VF_0208*'-gfp+ | This work |
| pKV496 | pEVS79 encoding FLP recombinase (KanR) | 69 |
| pVSV102 | Constitutive GFP for squid colonization assays | 75 |
| pEVS104 | Helper plasmid for conjugations | 59 |
| pEVS79-erm | Vector for allelic exchange (ErmR) | This work |
| pEVS79 | Vector for allelic exchange (CamR) | 59 |
| pMarVF1 | *V. fischeri* Mariner transposon vector (AmpR backbone, ErmR transposon) | 68 |

### DNA synthesis and sequencing

Primers used in this work are provided in **Table 3** and were synthesized by Integrated DNA Technologies (Coralville, IA). PCR products of gene deletions and plasmid modifications were validated using Sanger sequencing at Functional Biosciences (Madison, WI). Sequencing results were analyzed using Benchling. All Sanger sequencing products were generated with Q5 High-Fidelity polymerase (NEB), Phusion Hot Start Flex DNA polymerase (NEB), or OneTaq DNA polymerase (NEB). Diagnostic PCRs were performed with OneTaq DNA polymerase (NEB) or GoTaq DNA polymerase (Promega). Whole plasmid sequencing was performed by Plasmidsaurus (Eugene, OR) when noted.

**Table 3.**
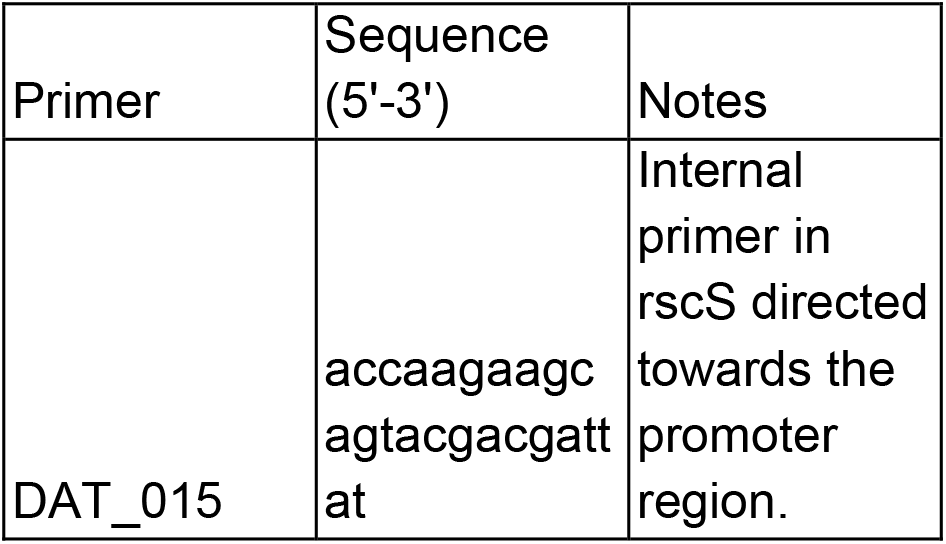
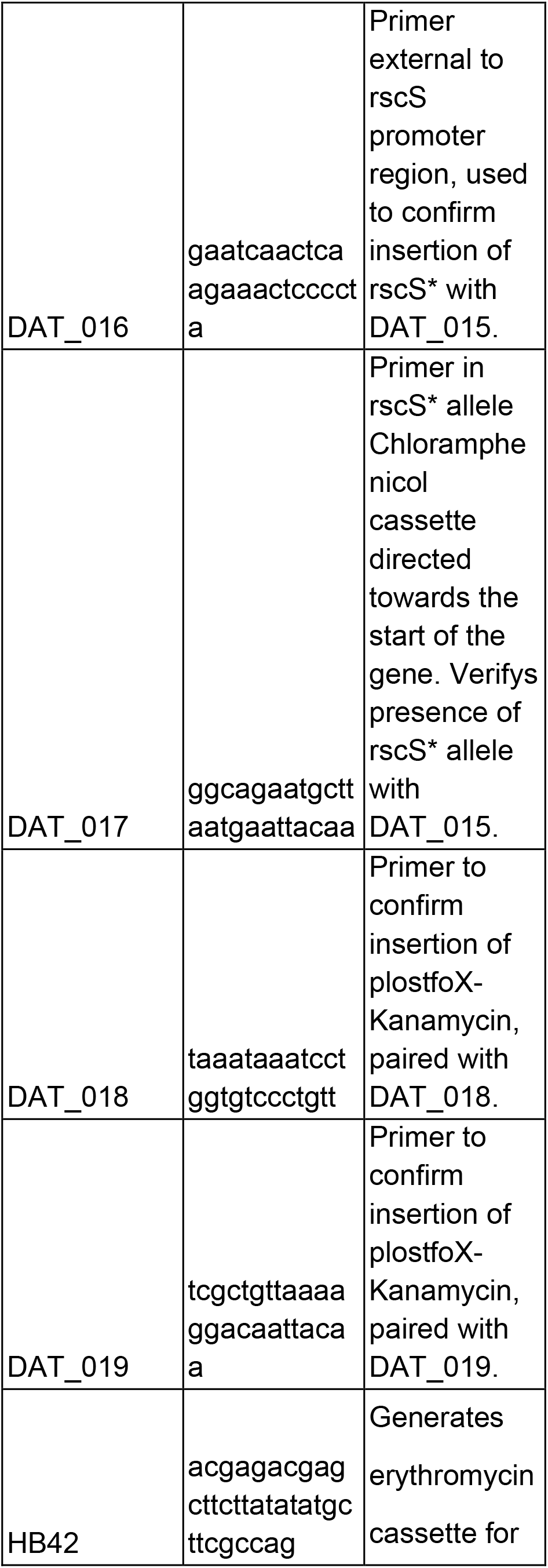
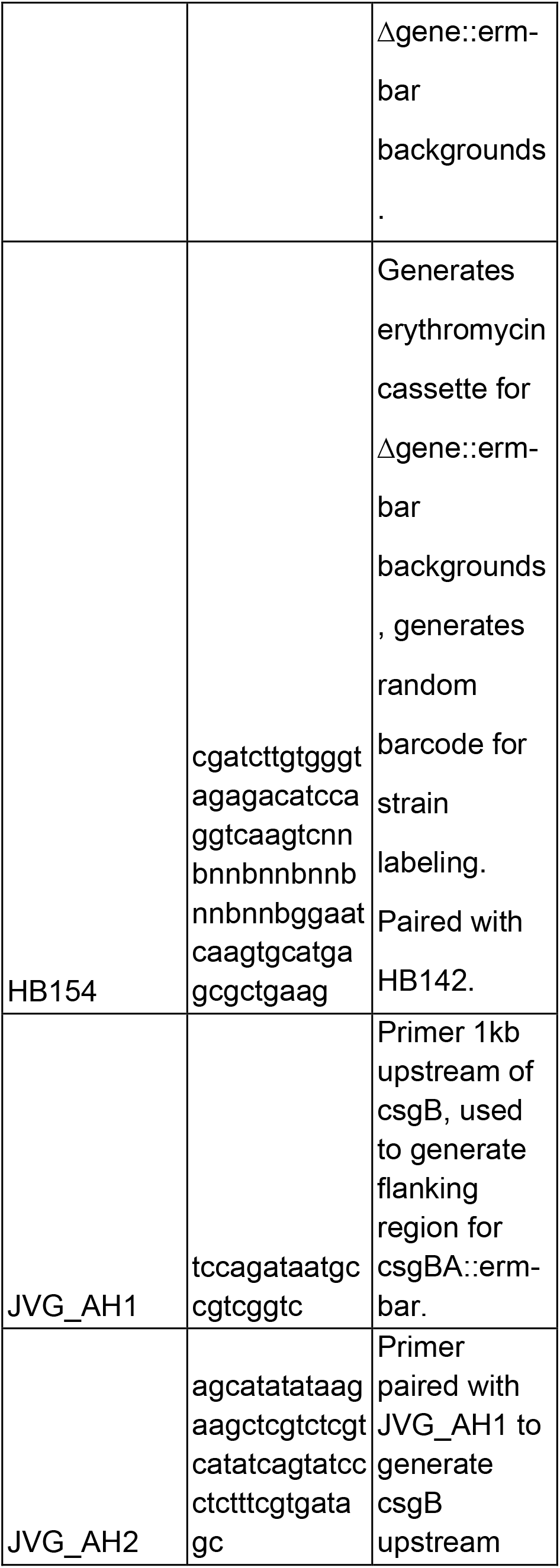
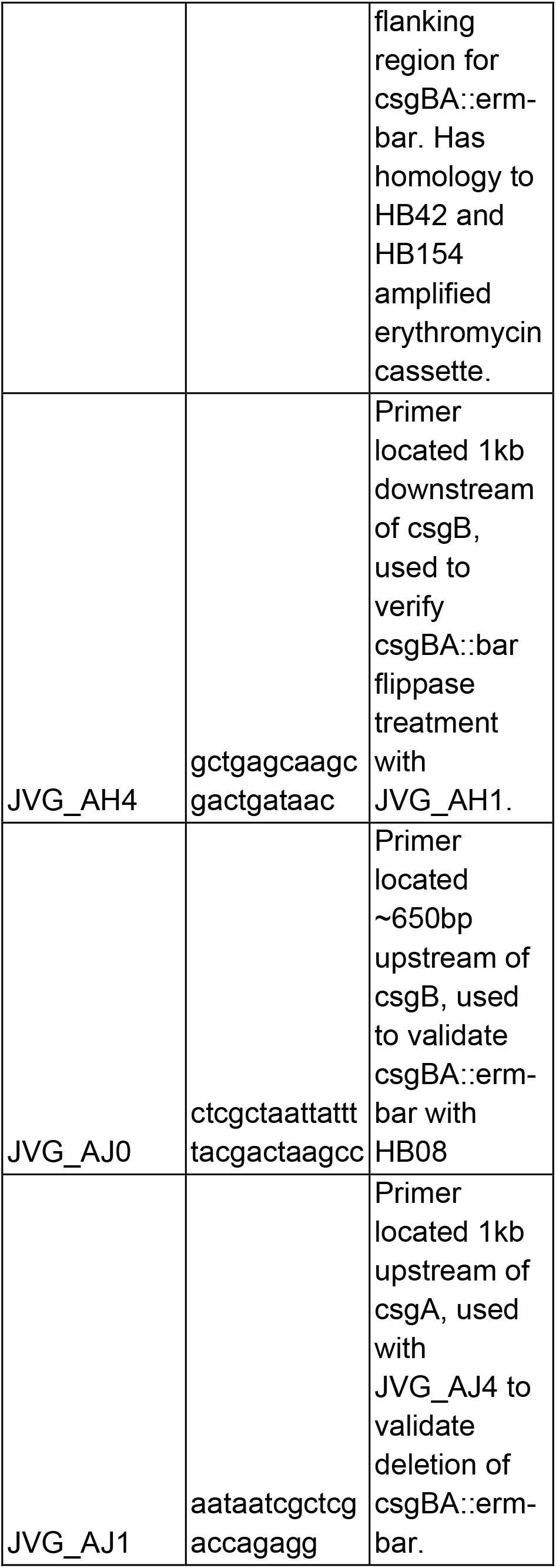
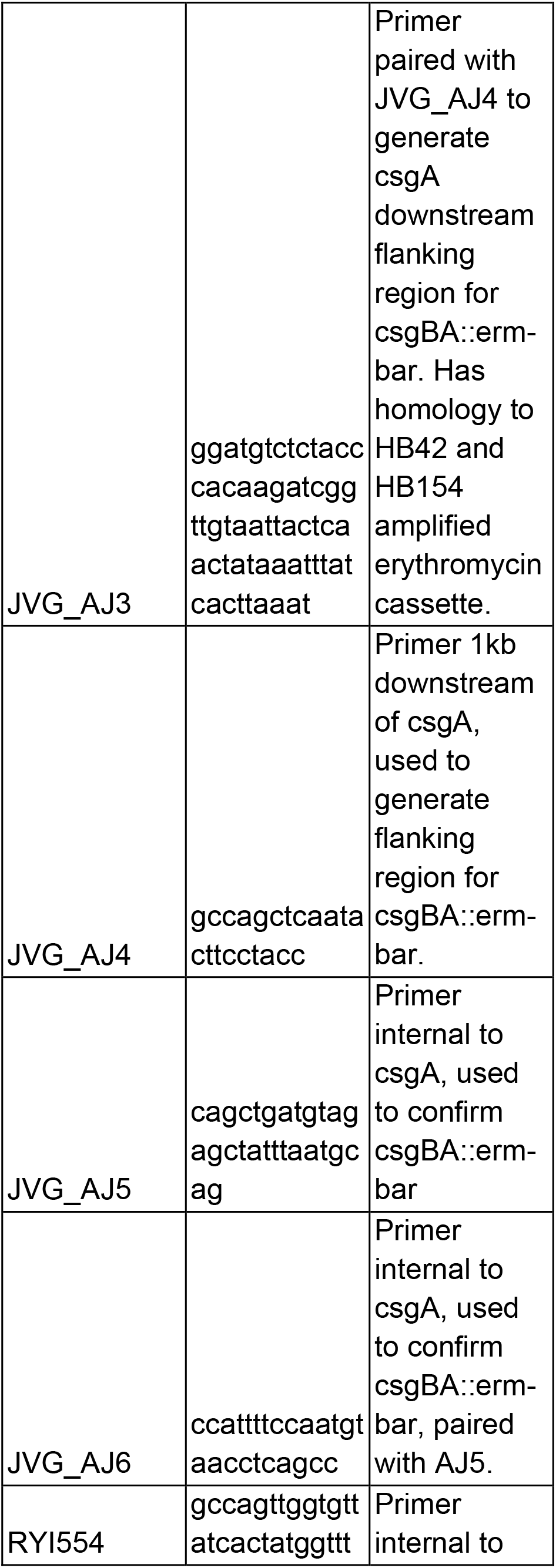
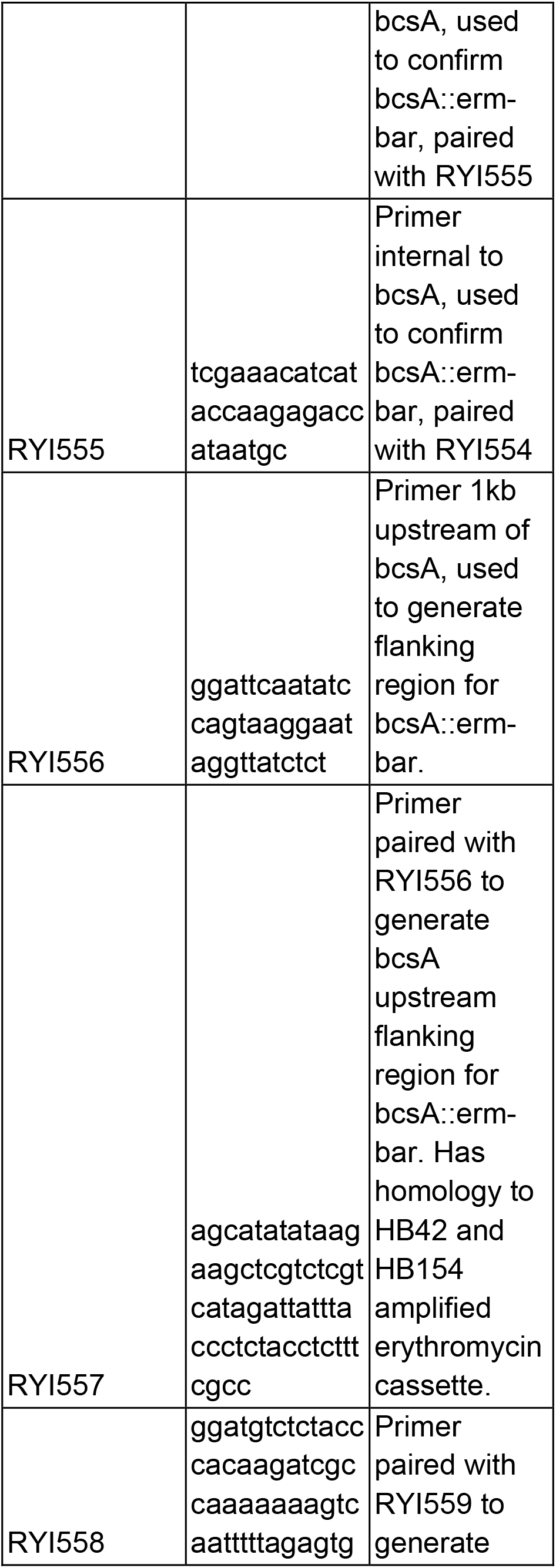
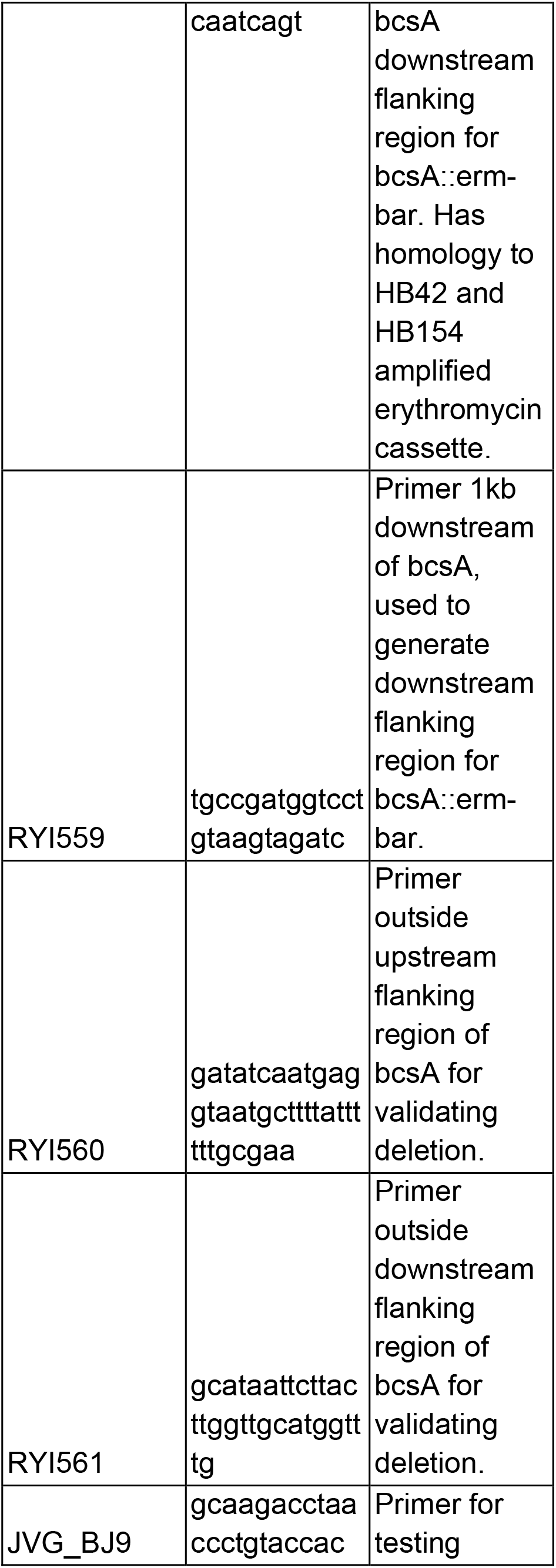
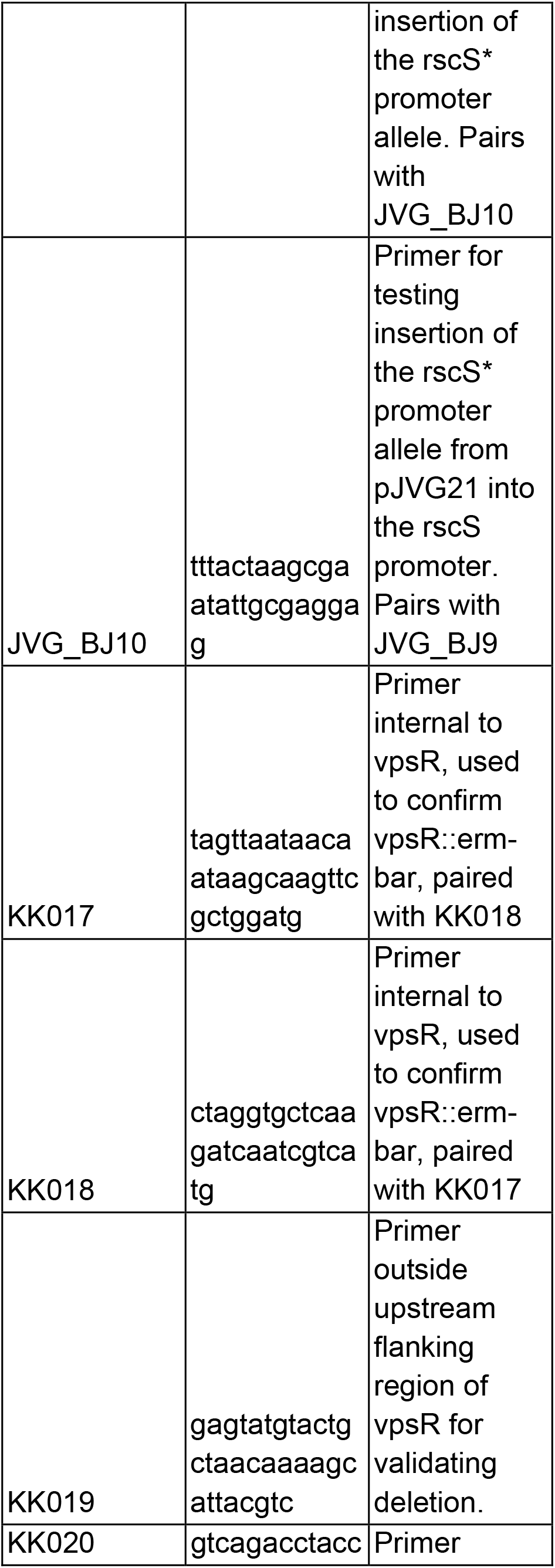
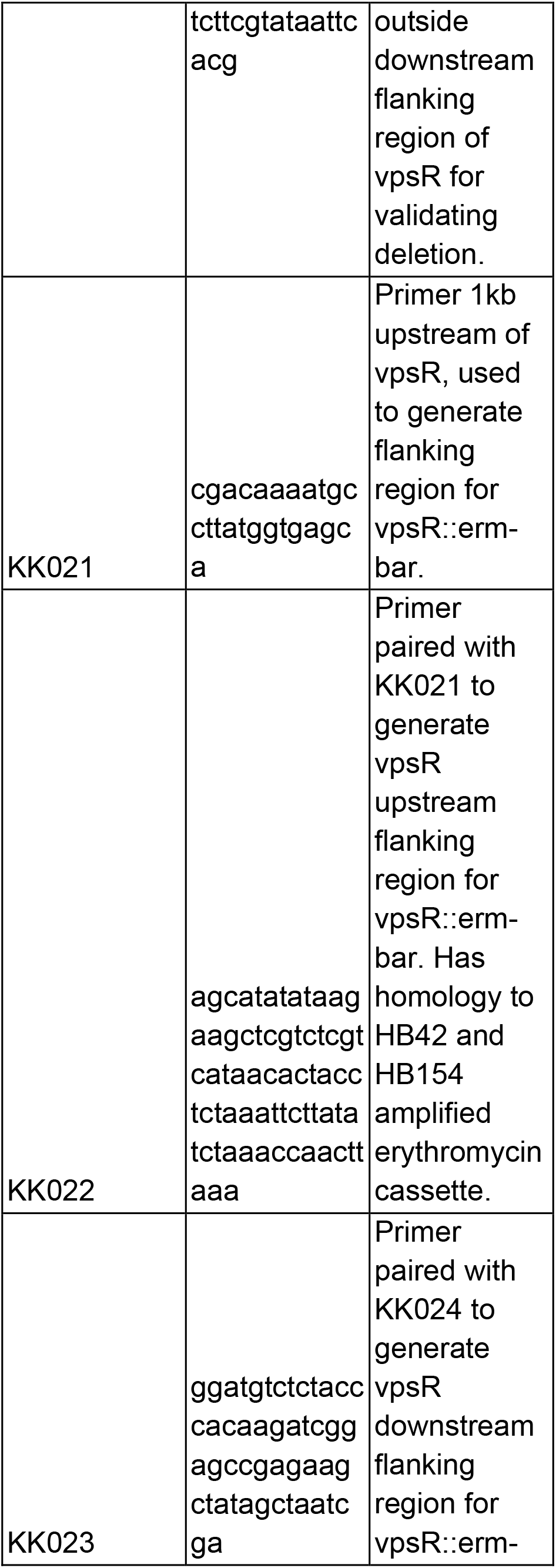
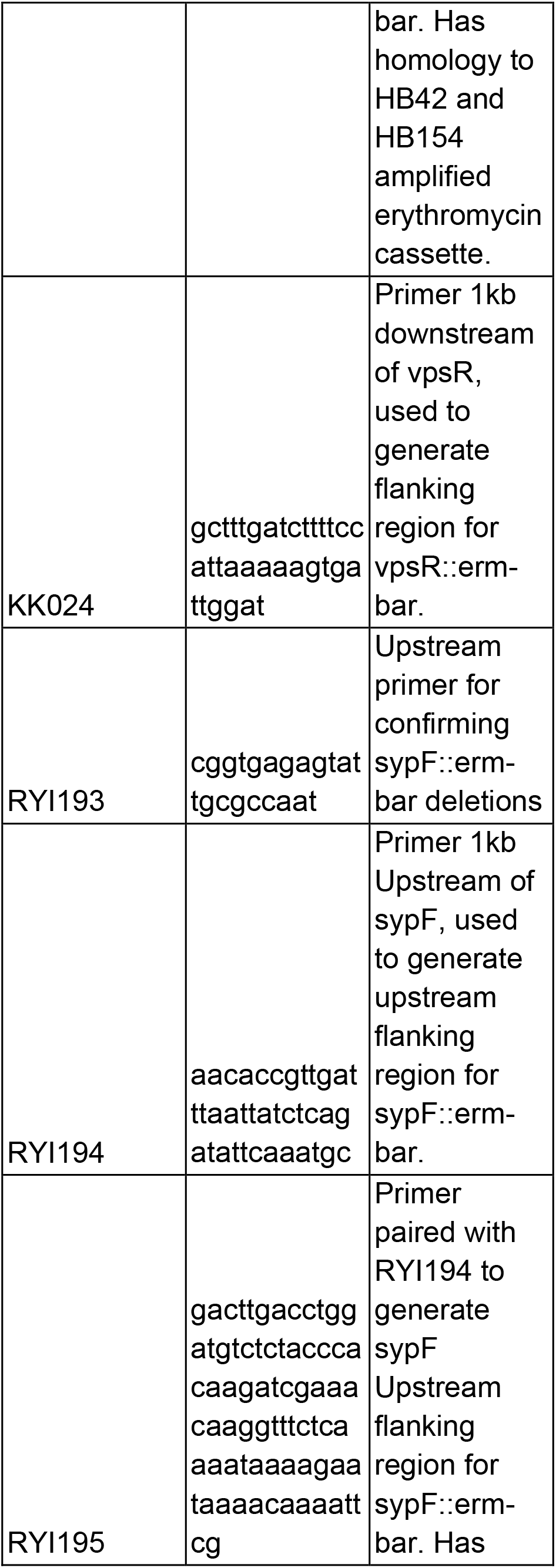
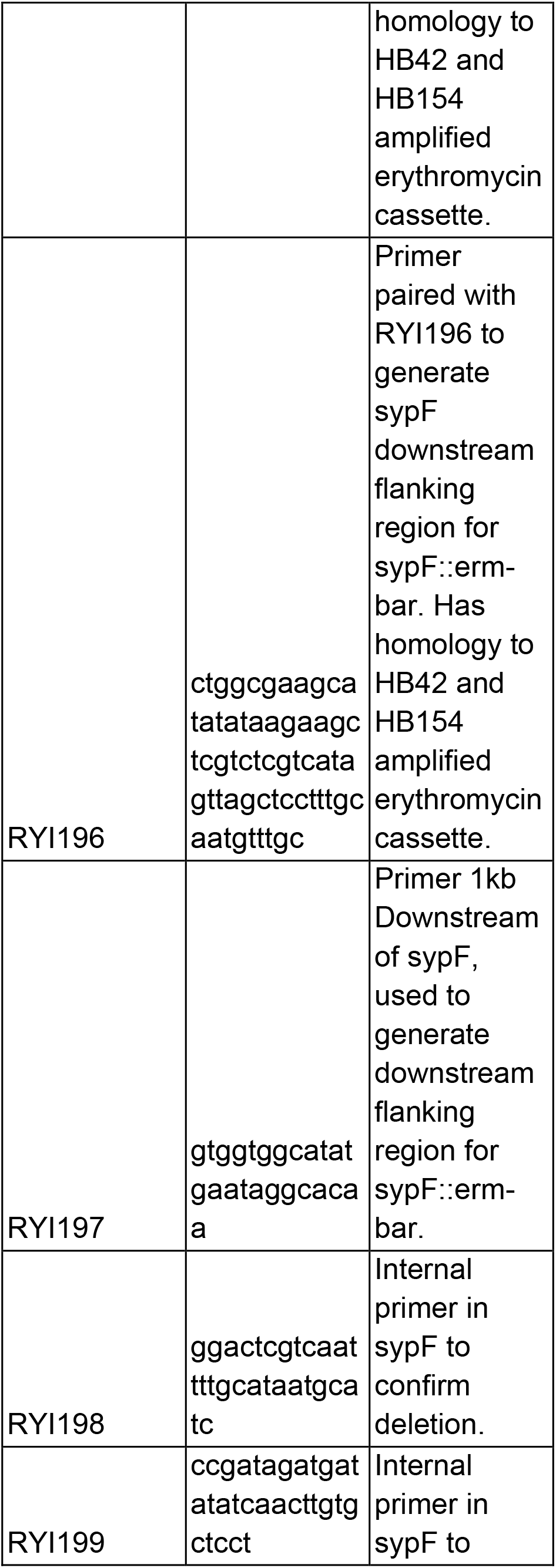
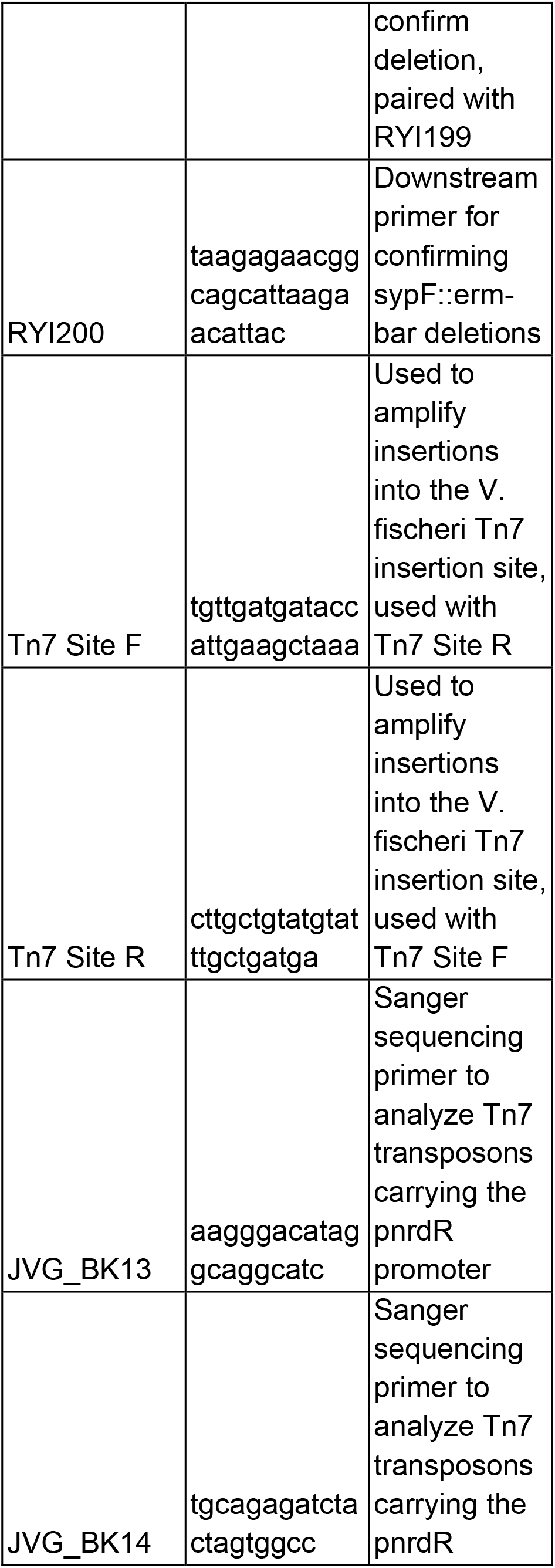
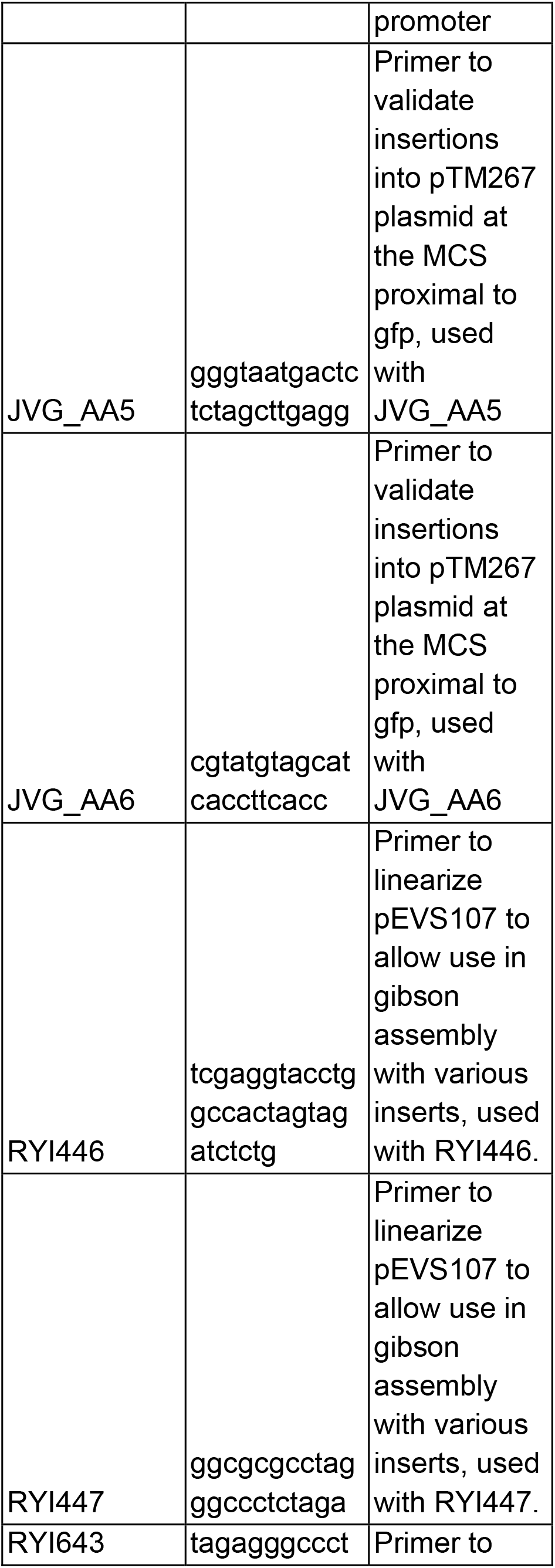
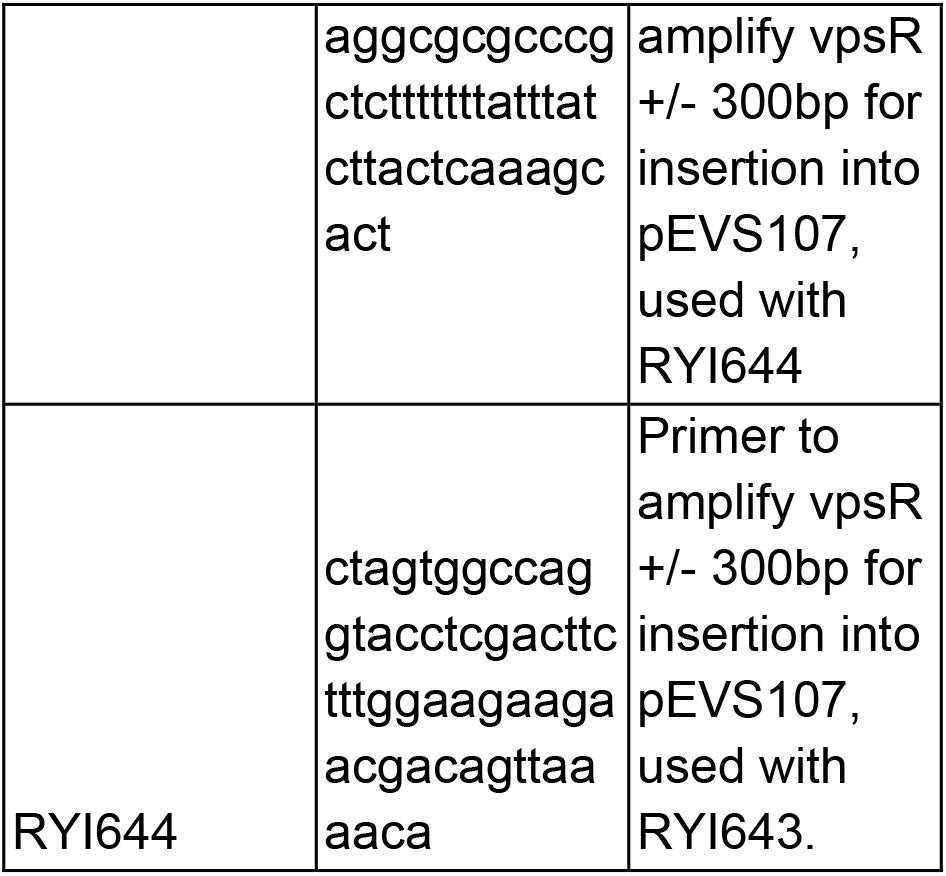
Primers:

### RNA isolation from *V. fischeri* wrinkled colonies

Strains MJM1100, MJM1198, MJM2251, and MJM3295 were streaked on LBS agar and grown overnight at 25 °C (30, 36, 60, 61). Overnight cultures were prepared in triplicate from independent colonies of each strain in LBS media, and grown overnight at 25 °C with rotation. From each overnight culture, 500µL was spotted onto LBS agar, or LBS-Ca^2+^ agar, and spots were allowed to grow for 48 hours at 25 °C. After 48 hours, each spot was collected into a 1:2 mix of LBS and RNAprotect Bacteria reagent (Qiagen), incubated at RT for 5 minutes, followed by vortexing at maximum speed on a Vortex Genie-2 (Scientific Industries), followed by centrifugation for 10 minutes at 5,000 xg at room temperature. Supernatants were removed, and cell pellets were stored at -80 °C. RNA extraction was performed using an RNeasy PowerBiofilm kit (Qiagen), as per manufacturer instructions. Purified RNA was stored at -80 °C, followed by a further on-column DNAse treatment using RNAse-free DNAse (Qiagen) followed by purification with the RNeasy MinElute kit (Qiagen). Samples were then transferred to the University of Wisconsin - Madison Biotechnology center.

### Library preparation, sequencing, and analysis of RNA samples

Samples were quality-tested using both NanoDrop and Agilent Bioanalyzer measurements. Ribosomal RNA depletion with additional probes recommended by Illumina (**Table 4**), stranded library preparation (Illumina Ribo-Zero Plus rRNA Depletion w/ Stranded Total RNA), and paired-end 2×150 sequencing was conducted at the UW-Madison Biotechnology center on an Illumina NovaSeq 6000. **Table S5** shows quality control data for these samples. The resulting reads were processed by FASTP v0.20.1 (62), mapped with BWA-MEM v0.7.17 (63), read counts per gene were enumerated with HTSeq v0.13.5 (64) to *V. fischeri* ES114 Chromosome 1, 2, and the natural plasmid pES100 (CP000020.2, CP000021.2, and CP000022.1), and differential expression analysis was conducted with EdgeR v3.32.1 (65). Follow-up analysis of the dataset was performed in RStudio v1.3.959. Volcano plots of data were produced through plotting the -Log_10_ (FDR) on the graph Y-axis and the Log_2_(Fold Change) on the X-axis, from the EdgeR results. Overlay graphs of multiple differential expression analyses were generated by plotting the Log_2_(Fold Change) for each gene in the stated differential expression analyses on either the X- or Y-axes. Transcripts per Kilobase Million (TPM) values for all genes were obtained from the raw read counts for all genes from HT-Seq output (39). The read counts for each gene were then divided by each gene’s respective length in kilobase pairs to generate reads per kilobase (RPK). These values were summed for each of the replicates and divided by 1,000,000 to generate the scaling factor for each replicate. The RPK for each gene was then divided by its respective replicate scaling factor to generate per gene TPM values. TPM values for all conditions can be found in **Table S6**

**Table 4.**
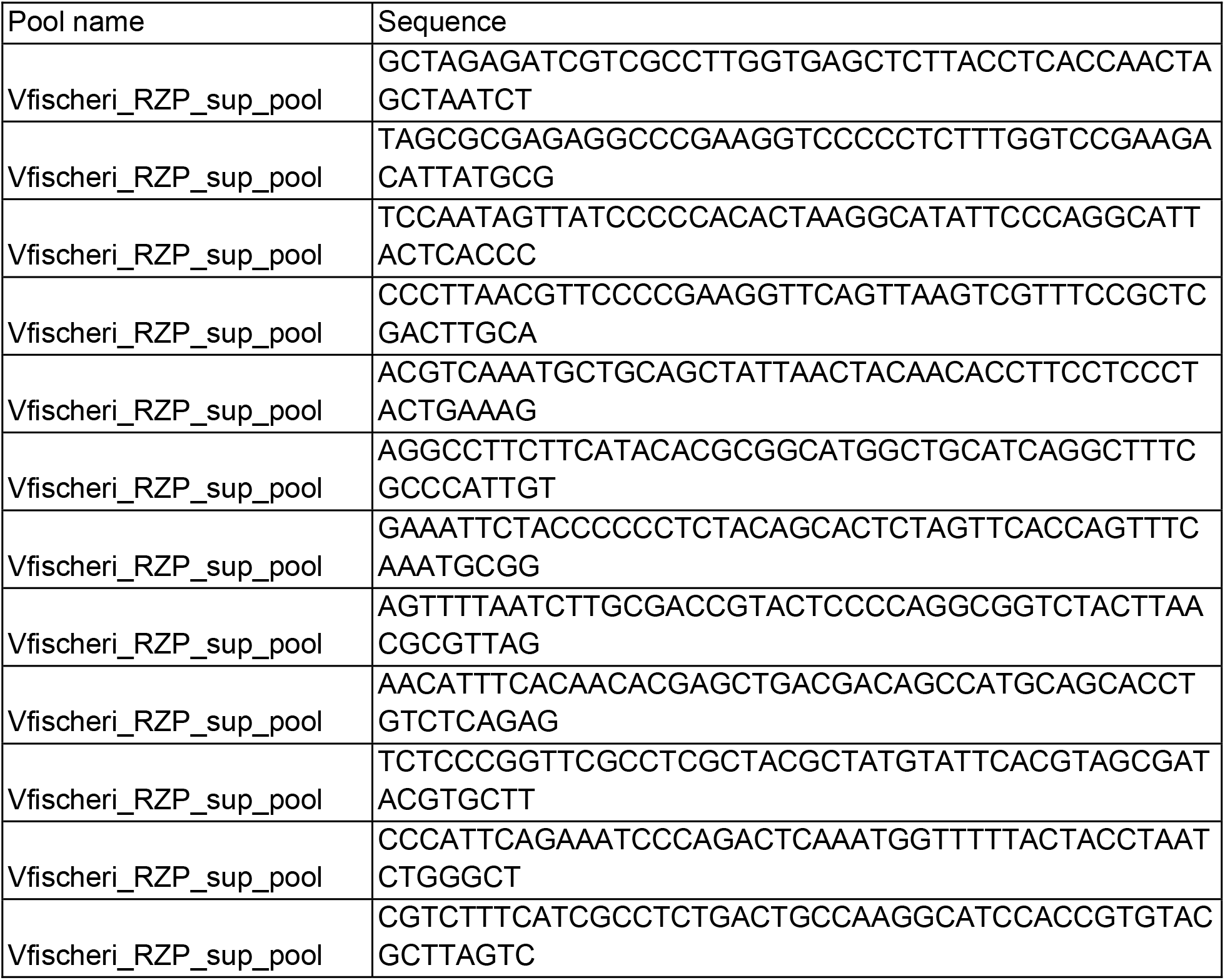
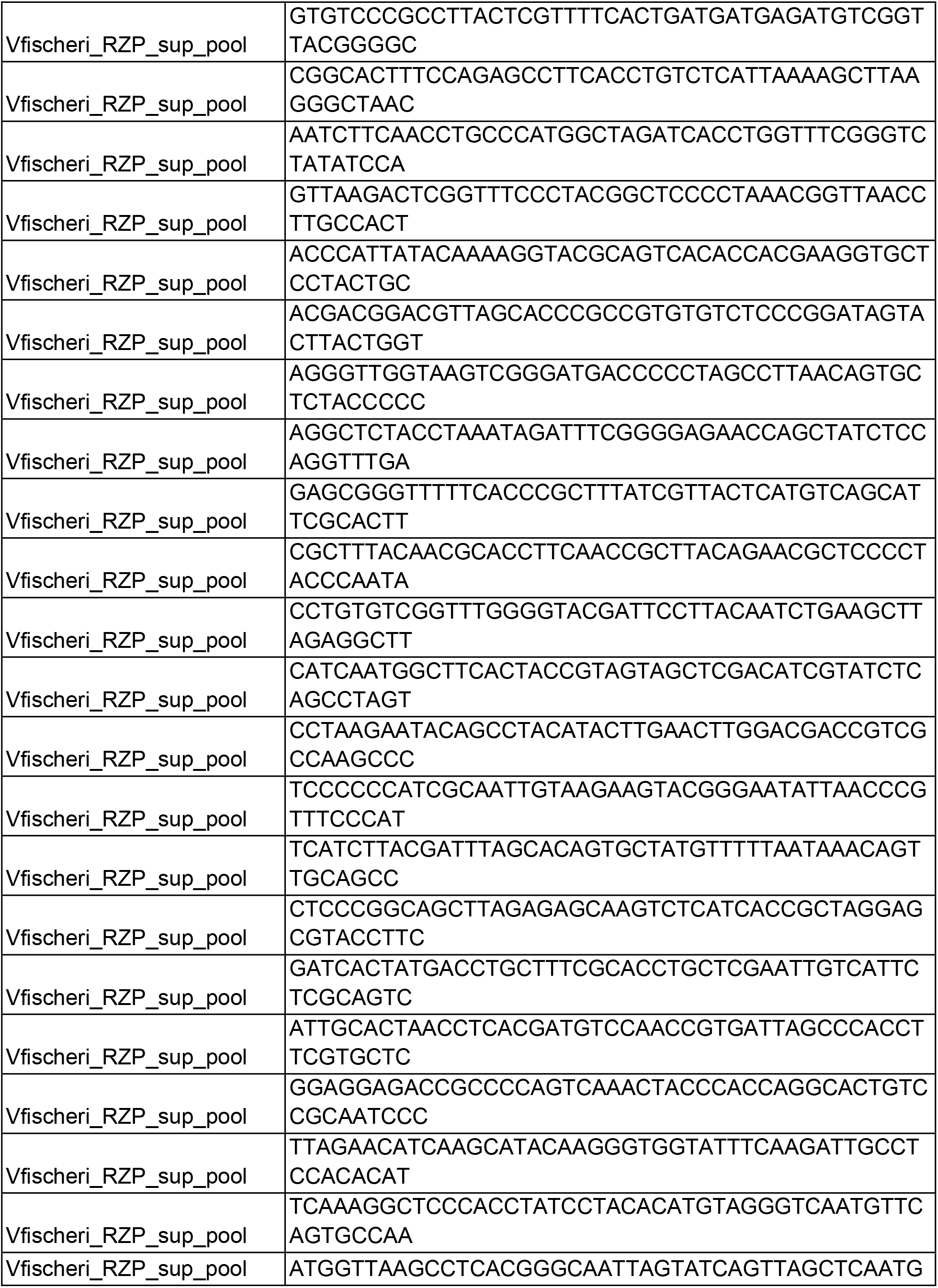
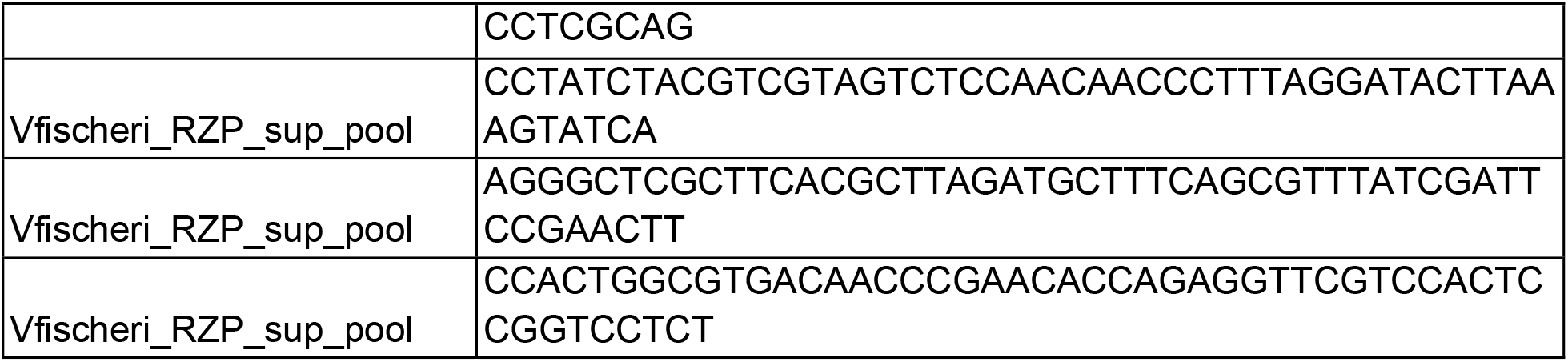
Additional Ribo Depletion Oligos:

### Construction of pTM267-based transcriptional reporters (pJVG11 and pJVG25)

Reporter plasmids were created through insertion of gene specific promoter elements into the established transcriptional reporter plasmid pTM267 (66). In short, ∼500 bp of upstream DNA preceding the start codon for *bcsE* and *VF_0208* were amplified using Q5 DNA Polymerase (NEB) using primers JVG_BC1 and JVG_BC2 (*bcsE*) and JVG_BM1 and JVG_BM2 (*VF_0208*).

Each primer set appended XbaI and XmaI restriction sites to promoter amplicons. Inserts and the pTM267 backbone were then digested with XbaI and XmaI (NEB), gel purified using the QIAquick gel extraction kit (QIAgen), and further concentrated using a DNA Clean and Concentrator kit (Zymo Research). Purified inserts and pTM267 backbone were then ligated together using overnight ligation at 16 °C with T4 DNA ligase (NEB). Ligation products were then transformed into chemically competent *E. coli* DH5ɑ λpir, and chloramphenicol resistant colonies were selected from plates. Potential candidates were streak purified, and used in colony PCR using primers JVG_AA5 and JVG_AA6 to amplify across the inserted element. Candidates showing the proper size amplicons were miniprepped and either sent for Sanger sequencing using primers JVG_AA5 and JVG_AA6 (pJVG11-*bcsE*) (Functional Biosciences) or for full plasmid sequencing (pJVG25-*VF_0208*) (Plasmidsaurus).

### Construction of *gene*::bar tagged gene deletions

The creation of Δ*gene*::bar mutants followed previously published protocols (67). In short, 1 kb regions flanking the gene(s) of interest were amplified from MJM1100 genomic DNA using Phusion Hot Start Flex polymerase (Thermo Fisher), and each flanking element shared terminal homology with a erythromycin resistance cassette amplified from plasmid pHB01 using primers HB42 and HB154. Splicing by Overlap Extension PCR (SOE-PCR) was then used to join the flanking regions onto the Erm^R^ cassette (67). This joined fragment was used as mutagenic DNA in a natural transformation into *V. fischeri* strains containing the pLostfoX-Chloramphenicol or pLostfoX-Kanamycin plasmid (68). Recombinants were selected for erythromycin resistance, streak purified, and PCR assays were used to identify candidates with insertions of the erythromycin resistance cassette in the place of targeted genes. For candidates passing PCR screens, the gene deletion and adjacent upstream and downstream regions were PCR amplified, and amplicons were sent for Sanger sequencing using primers HB08, HB09, HB42, and HB146. Candidates which successfully replaced the target gene with the erythromycin cassette were then saved as Δ*gene*::erm-bar strains. In order to eliminate potential influences of the erythromycin cassette on neighboring genes, the flippase-encoding plasmid pKV496 was conjugated into Δ*gene*::erm-bar strains via tri-parental mating with strains MJM534 and MJM3478 (69). Following this conjugation, candidates were validated for loss of the erythromycin cassette using PCR with primers flanking the deletion site, and amplicons were sent for Sanger sequencing for final validation. Candidates passing the deletion check were then saved as Δ*gene*::bar strains.

### Natural transformation of *V. fischeri* with genomic DNA

When necessary, plasmids pLostfoX-Chloramphenicol or pLostfoX-Kanamycin were conjugated into *V. fischeri* strains to induce competence to transfer alleles between strains (68, 70). Natural transformation was conducted as described previously (67), in which strains carrying pLostfoX- Chloramphenicol or pLostfoX-Kanamycin were grown from glycerol stocks in LBS with their respective antibiotics. After overnight growth in LBS, a 1/100 subculture was performed into Tris minimal media (TMM) containing the antibiotic of note, with further growth overnight. The following morning, the TMM overnight culture was subcultured 1/20 into fresh TMM and allowed to grow to OD600 0.2. This culture was then separated into 500 uL aliquots, and 2.4 µg genomic DNA was then provided to the aliquots as needed, and the cell/DNA mixtures were gently vortexed before resting at room temperature for 30 minutes. Following room temperature incubation, each aliquot was provided with 1 mL of LBS, and allowed to recover overnight at 25 °C. Recovery cultures were then plated on LBS agar containing respective antibiotics, and candidates of note were streak purified for downstream confirmation.

### Construction of Tn7 transposon-based chromosomal complementations of *sypF, sypG,* and *vpsR*

Chromosomal complementations of Δ*gene*::bar mutations were conducted through provisioning of the gene of interest on a Tn7 transposon. For the Δ*vpsR*::bar strain, *vpsR* +/- ∼350 bp was amplified from MJM1100 gDNA using primers RYI643 and RYI644, and the Tn7 transposon vector pEVS107 was linearized using primers RYI446 and RYI447 with Q5 DNA polymerase (NEB) (71). Using the NEB HiFi DNA assembly kit, the *vpsR* insert and pEVS107 backbone were joined, and the products of this reaction were transformed into *E. coli* DH5ɑ λpir, and erythromycin resistant colonies were selected. Candidates were streak purified, miniprepped using the QIAprep spin miniprep kit (Qiagen), and submitted to Plasmidsaurus for full plasmid sequencing, yielding plasmid pKK01. For the Δ*sypF* and Δ*sypG* complementations, the ORFs for each gene were amplified with JVG_BK7 and JVG_BK8 (*sypF*) and JVG_BK11 and JVG_BK12 (*sypG*) with Q5 DNA polymerase (NEB). A variant of pEVS107 containing the constitutively active promoter for the *nrdR* gene (pKMB036) was linearized using primers containing homology for *sypF* (JVG_BK5 and JVG_BK6) and *sypG* (JVG_BK9 and JVG_BK10), and the plasmid backbone and inserts were joined by the NEB HiFi DNA assembly kit.

Candidates were selected as erythromycin resistant colonies, and streak purified. Successful assemblies were confirmed using primers JVG_BK13 and JVG_BK14 across the insertion site using colony PCR, miniprepped using the QIAprep spin miniprep kit (Qiagen), and purified plasmids were sent to Plasmidsaurus for full plasmid sequencing, yielding plasmids pJVG22 and pJVG23.

### Chromosomal insertion of Tn7-complementations of *sypF, sypG,* and *vpsR*

The Tn7 transposon complementation plasmids of *sypF* (pJVG22), *sypG* (pJVG23), and *vpsR* (pKK01) were inserted into their respective *V. fischeri* Δ*gene*::bar deletion backgrounds using standard conjugation protocols, with the addition of the Tn7 transposase helper strain MJM637, carrying plasmid pUX-BF13 (59, 72). Candidates were selected as erythromycin resistant colonies, streak purified, and PCR confirmed using primers Tn7 Site F and Tn7 Site R with Phusion HS DNA polymerase. Tn7 Site F/R amplicons were then purified using the QIAquick PCR purification kit, and sent for Sanger sequencing using primers Tn7 Site F, Tn7 Site R, RYI460, and RYI461 (*vpsR*), or Tn7 Site F, Tn7 Site R, JVG_BK13, and JVG_BK14 (*sypF* and *sypG*) (Functional Biosciences). Passing candidates were then saved as MJM5035 (*rscS\** Δ*vpsR*::bar attTn7::*vpsR*-erm), MJM5303 (*rscS\** Δ*sypG*::bar attTn7::P*_nrdR_*-*sypG*-erm), and MJM5304 (*rscS\** Δ*sypF*::bar attTn7::P*_nrdR_*-*sypF*-erm)

### Construction of *rscS\** allelic exchange plasmid (pJVG21)

Plasmid pEVS79 (59) modified with ErmR replacing CamR (pEVS79-Erm, Matthew Hauserman) was amplified using primers JVG_BJ1 and JVG_BJ2, and the intergenic region containing the *rscS\** allele was amplified from strain MJM1198 using primers JVG_BJ3 and JVG_BJ4.

Products of the JVG_BJ1 and JVG_BJ2 reaction were digested prior to assembly using Dpn1 (NEB). The NEBuilder HiFi DNA Assembly kit (NEB) was then used to assemble JVG_BJ1/2 and JVG_BJ3/4, products of this reaction were then transformed into *E. coli* DH5α using heat-shock and transformants were selected on BHI-Erm150 media. Candidates were screened using PCR with primers JVG_BJ5 and JVG_BJ6, passing candidates were then submitted to Plasmidsaurus (Eugene, OR) for whole plasmid sequencing yielding plasmid pJVG21.

### Allelic exchange of rscS* into ΔsypF, ΔvpsR, ΔbcsA, and ΔcsgBA ΔbcsA

Plasmid pJVG21 was conjugated into MJM3972 (ES114 Δ*sypF*), MJM4549 (ES114 Δ*bcsA*), MJM4683 (ES114 Δ*vpsR*), and MJM4915 (Δ*bcsA,* Δ*csgBA*) using standard laboratory protocols. Conjugation spots were scrapped from plates, resuspended in 70 % IO, and plated on LBS-Erm^5^ plates to select for plasmid integration into the host genome. Candidates were streaked on LBS-Erm to purity, and individual colonies were patched on LBS-Erm, LBS-Cam, and LBS. Candidates showing Erm^R^, Cam^R^, and growth on LBS were then cultured in LBS-Erm^5^ and saved as single recombinants. Double recombinants were then generated as follows.

Single recombinants were first cultured overnight in LBS, overnight cultures were then diluted by 1/100 into fresh LBS and allowed to grow until OD600 reached 0.2. At this point, a further subculture was prepared by repeating the 1/100 dilution into LBS, and allowing the culture to grow once again to OD600 0.2. These subcultures were then plated on LBS media, and allowed to grow overnight at 25 °C. Resulting colonies were patched on LBS-Erm, LBS-Cam, and LBS, and grown overnight at 25 °C. Patches showing Erm^S^ dropped the plasmid backbone, yet retained the Cam^R^ of the *rscS*\* allele were then cultured and saved as candidate double recombinants. To confirm insertion of *rscS\**, check PCRs were conducted using primers DAT_015 and DAT_016 and Phusion HS polymerase. Amplicons showing the proper size of the *rscS\** insertion were purified using the QIAgen Quickspin PCR purification kit, and sent to Functional Biosciences for Sanger sequencing confirmation. Passing candidates were saved as MJM4917 (*rscS\** Δ*sypF*), MJM4918 (*rscS\** Δ*bcsA*), MJM4919 (*rscS\** Δ*vpsR*), and MJM4956 (*rscS\** Δ*bcsA* Δ*csgBA*).

### Wrinkled colony assays

Bacterial strains were grown in LBS medium for approximately 17 hours, and 8 µL of each overnight culture was then spotted onto LBS or LBS-Ca^2+^ plates, and incubated at 25 °C. At 24 hours and 48 hours post spotting, plates were imaged using a Leica M60 microscope and Leica DFC295 camera.

### Plasmid-based expression assay

Bacterial strains were grown in LBS-Cam medium for approximately 17 hours at 25 °C, and 8 µL of each overnight culture was then spotted onto LBS-Cam and LBS-Cam-10mM Ca^2+^ Omnitrays, followed by incubation at 25 °C. After 24 hours, colony spots were imaged on a Zeiss Axio Zoom.v16 large-field 561 fluorescent stereo microscope, and fluorescence levels were measured within colony spots using the Zen Blue software. Expression of the reporter (eGFP fluorescence, 488 nm excitation, 509 nm emission, 300 ms exposure time) was normalized to a constitutive locus on the pTM267 backbone (mCherry fluorescence, 558 nm excitation, 583 nm emission, 700 ms exposure time) as eGFP fluorescence/mCherry fluorescence. Data analysis was conducted in GraphPad Prism using an ordinary one-way ANOVA with multiple comparisons.

### Soft agar swimming motility assay

Bacterial strains were streaked on TBS-agar 48 hours prior to the assay, and incubated at 25 °C. After 24 hours, four single colonies were cultured into TBS medium, and grown at 25 °C for 17 hours with rotation. 10uL of each overnight culture was then spotted onto TBS motility media (3g/L agar) in technical triplicate, and allowed to incubate at 25 °C. After 5 hours, plates were imaged using a Nikon D810 digital camera, and ImageJ was used to quantify the Fercart diameter of each migration spot as the migration distance. Each biological replicate was produced through averaging the technical replicates. Data analysis was conducted in GraphPad Prism using an ordinary one-way ANOVA with multiple comparisons. This assay was conducted on two separate days, for a total of 8 biological replicates.

### Congo red assay

Bacterial strains were grown in LBS medium for 17 hours, and 8 uL of each overnight culture was spotted on LBS Congo red media. Plates were incubated overnight at 25 °C for 24 hours, after which colony spots were transferred to printer paper, air dried, and imaged as TIFF files (69).

### Cyclic-di-GMP reporter assay

Bacterial strains were grown in LBS-Gent at 25 °C for 16 hours, and 8 uL of each overnight culture was spotted in technical duplicate on LBS-Gent and LBS-Gent-Ca^2+^ plates. After 24 hours, colony spots were imaged on a Zeiss Axio Zoom.v16 large-field 561 fluorescent stereo microscope, and fluorescence levels were measured within colony spots using the Zen Blue software. Expression of the reporter (TurboRFP fluorescence, 553 nm excitation, 573 nm emission, 150 ms exposure time) was normalized to a constitutive locus on the pFY4535 backbone (AmCyan1 fluorescence, 467 nm excitation, 496 nm emission, 150 ms exposure time) as TurboRFP fluorescence/AmCyan1 fluorescence. Data analysis was conducted in GraphPad Prism using an ordinary one-way ANOVA with multiple comparisons. For between condition analysis of LBS vs LBS-Ca^2+^ data, data analysis was conducted in GraphPad Prism using a two-way ANOVA with Šídák’s multiple comparisons test.

### Curli mutant squid single strain colonization assay

Lab reared *E. scolopes* hatchlings were colonized using a previously published protocol (73). In short, 3-5 x 10^3^ CFUs of the wild-type strain MJM1100 or the Δ*csgBA* mutant strain MJM4185 were provided to squid hatchlings, followed by a three hour colonization period. After 48 hours, hatchlings were measured for luminescence using a Promega GloMax 20/20 luminometer and euthanized. Homogenization of squid tissues was carried out as in the protocol, and colonization levels were enumerated from CFU/mL plated on LBS agar. This experiment was conducted in biological triplicate. A Mann-Whitney test was then used to determine statistical significance between the colonization levels of MJM4185 and wild type in GraphPad Prism.

### Curli mutant squid aggregation assay

Squid aggregation experiments were performed as described previously (74). In short, cultures of MJM1107 and MJM4276 were grown overnight at 25 °C in LBS-Kan (75). The following morning,10^5^ to 10^6^ CFU/mL of each strain were added to bowls of *E. scolopes* hatchlings and allowed to colonize for 3 hours. Following this colonization period, hatchlings were anesthetized in 2 % Ethanol/FSIO for 5 minutes, before fixation in 4 % Paraformaldehyde at 4 °C for 48 hours. Following fixation, hatchlings were washed 4 times with 1X mPBS (50 mM Sodium Phosphate, 0.45 M NaCl [pH 7.4]) and dissected and imaged with a Zeiss Axio Zoom.v16 large-field 561 fluorescent stereo microscope. Aggregate sizes were determined using the aggregate area tool, and statistical analysis was conducted using the Mann-Whitney test on GraphPad Prism.

## DATA AVAILABILITY

Transcriptomic data were deposited in the NCBI GEO under Series GSE237189 (Accession numbers GSM7596716-GSM7596730).

## ACKNOWLEDGEMENTS

We thank Hector Burgos for strain MJM3381, Katie Bultman for plasmid pKMB036, and Matthew Hauserman for plasmid pEVS79-Erm. This study was funded by NIGMS grant R35 GM148385 to M.J.M., R.Y.I was supported by NIGMS training grant T32 GM007215, and J.A.V.G. was supported by NIAID training grant T32 AI007414.

